# Valorization of CO_2_ through lithoautotrophic production of sustainable chemicals in *Cupriavidus necator*

**DOI:** 10.1101/2020.02.08.940007

**Authors:** Shannon N. Nangle, Marika Ziesack, Sarabeth Buckley, Disha Trivedi, Daniel M. Loh, Daniel G. Nocera, Pamela A. Silver

**Affiliations:** Department of Systems Biology, Harvard Medical School, Boston, MA, USA; Wyss Institute for Biologically Inspired Engineering, Boston, MA, USA; Department of Earth and Environment, Boston University, Boston, MA, USA; Faculty of Arts and Sciences, Harvard University, Boston, MA, USA; Department of Chemistry, Harvard Medical School, Boston, MA, USA

## Abstract

Coupling recent advancements in genetic engineering of diverse microbes and gas-driven fermentation provides a path towards sustainable commodity chemical production. *Cupriavidus necator* H16 is a suitable species for this task because it effectively utilizes H_2_ and CO_2_ and is genetically tractable. Here, we demonstrate the versatility of *C. necator* for chemical production by engineering it to produce three products from CO_2_ under lithotrophic conditions: sucrose, polyhydroxyalkanoates (PHAs), and lipochitooligosaccharides (LCOs). We engineered sucrose production in a co-culture system with heterotrophic growth 30 times that of WT *C. necator*. We engineered PHA production (20-60% DCW) and selectively altered product composition by combining different thioesterases and phaCs to produce copolymers directly from CO_2_. And, we engineered *C. necator* to convert CO_2_ into the LCO, a plant growth enhancer, with titers of ∼1.4 mg/L—equivalent to yields in its native source, *Bradyrhizobium*. We applied the LCOs to germinating seeds as well as corn plants and observed increases in a variety of growth parameters. Taken together, these results expand our understanding of how a gas-utilizing bacteria can promote sustainable production.

## Introduction

A sustainable future relies, in part, on minimizing the use of petrochemicals and reducing greenhouse gas (GHG) emissions^1^. Modern society relies on fossil fuels for power, transportation, and chemical production but lacks clear paths towards viable substitutes. As industrial bioproduction has grown, economies of scale and use of cheaper feedstocks show promising trends towards commodities^2,3^. Some of the cheapest and most sustainable feedstocks are gases (e.g., CO, CO_2_, H_2_, CH_4_) from various point sources: e.g., steel mills, ethanol production plants, steam reforming plants, and biogas^4,5^. Compared to commonly used carbohydrate-based feedstocks, these gas sources deliver carbon and energy sources to microbes in gas fermentation, and have the potential to be more cost-effective, use land more efficiently, and have a smaller carbon footprint^2^. In the past decade, synthetic biology has developed a multitude of tools that enable the synthesis of complex molecules along with the means to domesticate non-model microbes. Use of these genetic tools for autotrophic strains could lead to widespread adoption of gas fermentation^6–8^ and advance bioproduction of commodities.

*C. necator* H16 (formerly known as *Ralstonia eutropha* H16) is among the most attractive species for industrial gas fermentation. It is a facultative Knallgas bacterium that can derive its energy from H_2_ and carbon from CO_2_, is genetically tractable, can be cultured with inexpensive minimal media components, is non-pathogenic, has a high-flux carbon storage pathway, and fixes the majority of fed CO_2_ into biomass^6–8^. Over the past few decades, *C. necator* has been used for polyhydroxyalkanoate (PHA) production with potential commercial production. In addition to PHAs, *C. necator* has been engineered to produce a variety of compounds^9^ including: 2-methylcitric acid^10^, ferulic acid^11^, isopropanol and 3-methyl-1-butanol^12^—under heterotrophic conditions. More recently, lithotrophic conversion of CO_2_ (via aerobic oxidation of hydrogen) by engineered strains has produced: 600 mg/L isopropanol, and 220 mg/L C_4_ / C_5_ fusel alcohols^13^, alkanes and alkenes at 4.4 mg/L^14^, methyl ketones at 180 mg/L^15^, and stable-isotope-labeled arginine at 7.1% of dry cell weight (DCW)^16^. We previously showed lithotrophic production of C_3_ and C_4_ / C_5_ alcohols in a hybrid biological-inorganic system that supplies CO_2_ and generates H_2_ from water-splitting electrodes directly in the culture medium^13^. Here, we expand on this previous work by engineering *C. necator* to produce a larger diversity of products that seek to promote the sustainable development of industrial bioproduction.

Our efforts use *C. necator* to bridge the gap between cheap gaseous feedstocks and versatile bioproduction. We address three avenues for bioproduction, selected for their potential to reduce GHG emissions if industrially scaled. First, if bioproduction is to play a major role in replacing unsustainable industries, we must attempt to provide for existing infrastructure by producing feedstocks for heterotrophs from CO_2_, rather than from plant material. Second, to demonstrate the versatility of commodity products *C. necator* is well-positioned to address, we sought to diversify the types of PHA copolymers that can be made lithotrophically—beyond P(3HB). Third, we use *C. necator* to produce a plant growth enhancer to promote crop yields and offset fertilizer use. Implementation of these three avenues can reduce the demands set on agriculture to generate bioproducts while increasing land-use efficiency for food.

## Results

### Sugar feedstock production

We engineered *C. necator* to convert CO_2_ into sucrose as feedstock for the heterotrophs *E. coli* and *S. cerevisiae* (for experimental setup see **Supplementary Fig. 1a**). Sucrose has high energy-density and is commonly used by other heterotrophs. To produce sucroses in *C. necator*, we engineered it to express the cyanobacterial sucrose synthesis enzymes and a sucrose porin to export the sucrose into the media. The engineered pathway is shown in **Fig. 1a**: sucrose phosphate synthase (SPS) catalyzes the conversion of UDP-glucose and fructose 6-phosphate into sucrose 6-phosphate and sucrose phosphate phosphatase (SPP) hydrolyzes the phosphate to yield sucrose.

**Figure 1.**
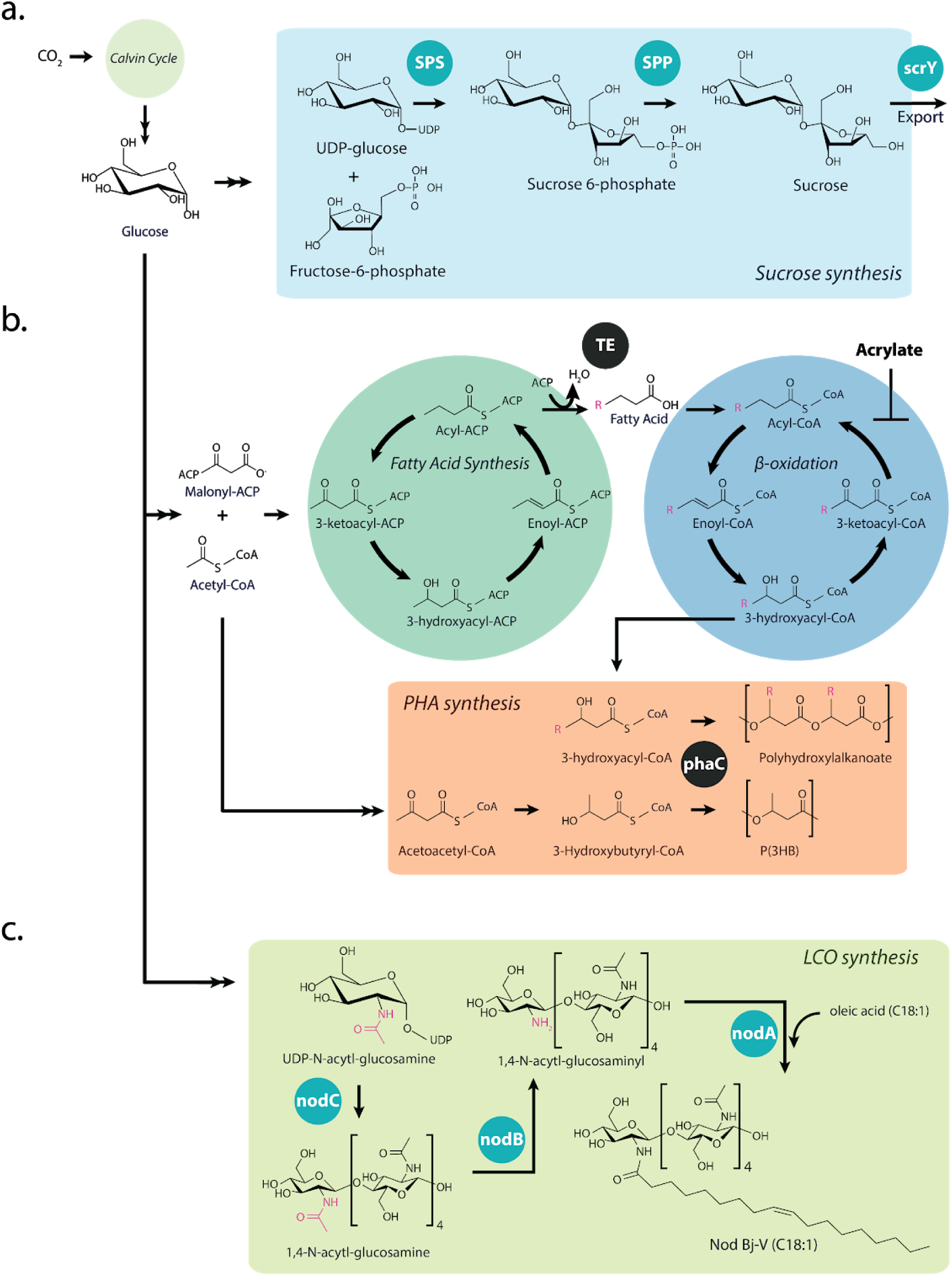
Metabolic pathways that were modified in *C. necator*. (**a**) Sucrose synthesis pathway. Enzymes from *Synechocystis sp.* PCC 6803: SPS, sucrose phosphate synthase, which binds UDP-glucose and fructose-6-phosphate and SPP, sucrose phosphate phosphatase, which removes the phosphate on the fructose to release sucrose. Enzymes from *E. coli*: scrY, sucrose porin, which exports sucrose through facilitated diffusion. (**b**) PHA synthesis pathway. Thioesterase (TE) enzymes from: *U. californica* FatB2 a 12:0 acyl-ACP TE and an engineered chimera of *C. palustris* FatB1(aa 1-218) and FatB2 (aa 219-316)—Chimera 4 (chim4), which produce free fatty acids of specific lengths. PHA synthase (phaC) enzymes: Native *C. necator* C4 phaC*_Cn_*; *P. aeruginosa PAO1* phaC2*_Pa_*; *P. aeruginosa PAO1* phaC1*_Pa_*; and a *Pseudomonas spp 61-3* phaC1*_Ps_*, which perform the final step in PHA polymerization and grow the chain. (**c**) Nodulation factor synthesis pathway for Nod Cn-V (C_18:1_). Enzymes from *B. japonicum*: NodC protein, an N-acetylglucosaminyltransferase that builds the backbone, NodB, a deacetylase that acts on the non-reducing end, and NodA, an acetyltransferase that attaches a fatty acid.

To choose the optimal enzymes for engineering, we compared the titers of overexpressed SPS and SPP from *Anabaena cylindrica* PCC 7122 and *Synechocystis sp.* PCC 6803, and a fusion SPS from *Synechococcus elongatus* PCC7942 in *C. necator*. We found the *Synechocystis* enzymes led to 10-fold higher titers compared to those from *A. cylindrica* (**Supplementary Fig. 2**). We did not detect any production from the *S. elongatus* enzymes. Thus, all subsequent experiments were performed in *C. necator* strains containing the *Synechocystis* sucrose synthesis enzymes. Expression of only SPS and SPP led to sucrose titers of about 100 mg/L (8 days post induction) (**Fig. 2a**). When we added the genes encoding the sucrose porin scrY from *E. coli*, we detected an 80% increase in titers in the supernatant, to ∼180 mg/L and cell viability post-induction improved (data not shown). The increases in titer from adding sucrose export indicates that actively removing it from the cytosol could increase enzymatic production by lowering the inhibitive effects of product accumulation.

**Figure 2:**
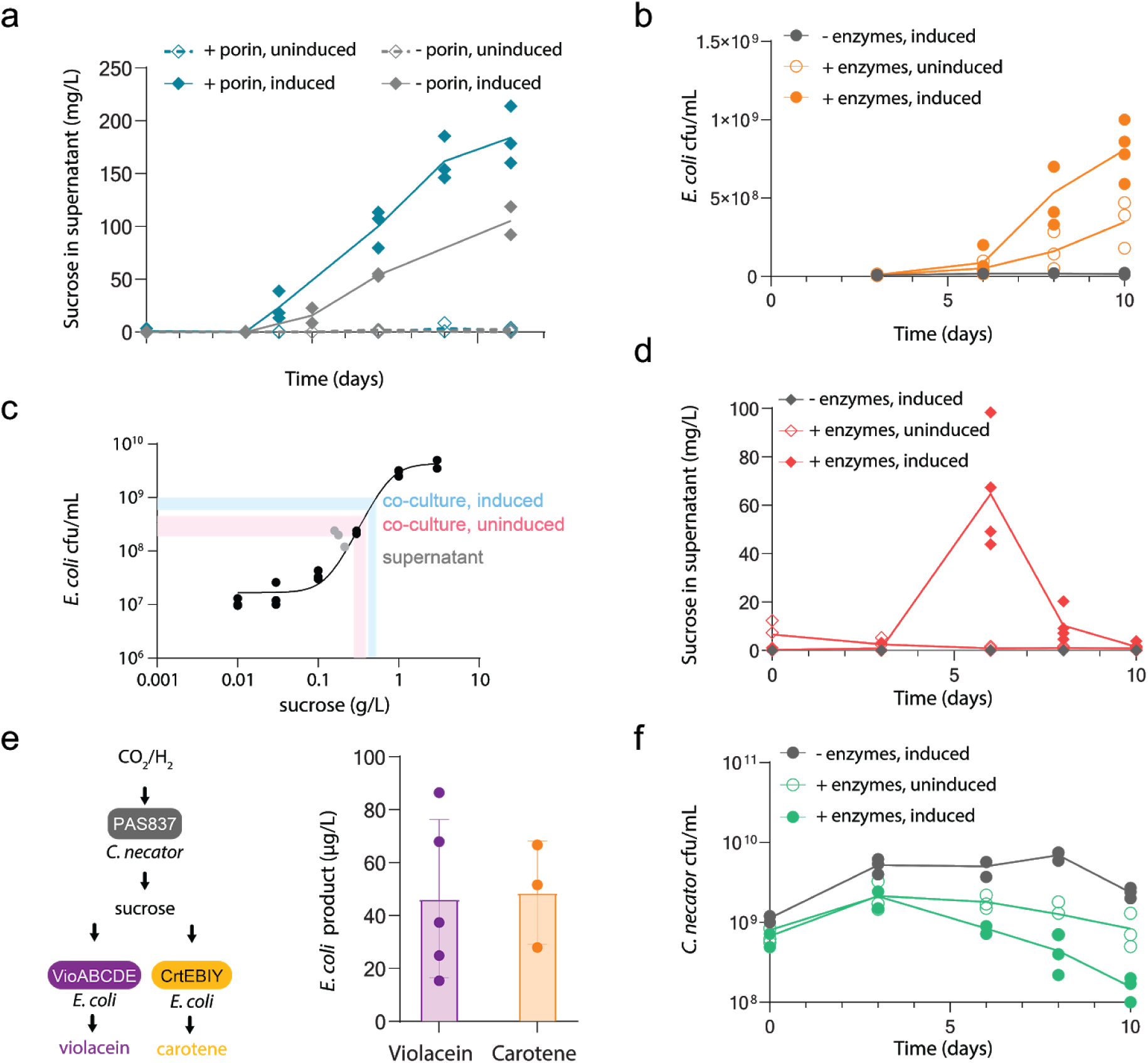
Sucrose-based *C. necator*-*E. coli* co-culture fueled by CO_2_/H_2_. Depicted are average values with error bars indicating standard deviation. (**a**) Sucrose production in supernatant from *C. necator* without porin (grey diamonds) or with porin (blue diamonds) without arabinose induction (empty symbols) or with 0.3% arabinose after 3 days (filled symbols). (**b**) *C. necator-E. coli* co-culture. PAS842 growth in co-culture with WT *C. necator* H16 and induction (grey circles), with PAS837 without induction (empty orange circles) or with induction (filled orange circles). (**c**) Comparison of PAS842 growth in supernatant derived from PAS837 with co-culture. PAS842 was grown in increasing sucrose concentrations to generate a standard curve. PAS842 grown in PAS837 supernatant in relation to measured sucrose in supernatant is indicated in grey circles. PAS842 growth in co-culture with uninduced PAS837 (empty orange circles) and induced PAS837 (filled orange circles) are shown. Ranges for uninduced and induced are 0.28-0.39 g/L and 0.42 - 0.53 g/L, respectively. (**d**) Sucrose concentrations in the supernatant of conditions as described above (grey diamonds, grey empty circles, grey filled circles respectively). (**e**) Violacein and carotene production in co-culture. PAS845 and PAS846 were grown in co-culture with PAS837 with induction. Samples were harvested, violacein and carotene were extracted and quantified as described in the Methods. (**f**) *C. necator* growth in conditions as described above (grey circles, empty green circles, filled green circles respectively).

We then demonstrated that the sucrose produced by *C. necator* can serve as a carbon source for heterotrophs. To test the heterotrophs’ growth response, we used engineered *E. coli* and *S. cerevisiae* strains with enhanced sucrose utilization: *E. coli* ΔcscR W (PAS842) and *S. cerevisiae* W303^Clump^ (PAS844). The *E. coli* strain (PAS842) was engineered with genomically-integrated sucrose catabolism (csc) operon containing an invertase (CscA), a sucrose permease (CscB), and a fructokinase (CscK); the repressor (CscR) was knocked out^17–19^. This strain was observed to grow to equivalent densities from ∼5x lower sucrose than the parent W strain (**Supplementary Fig. 3**). The yeast strain (PAS844) was derived from a low-sucrose directed evolution experiment that conferred increased sucrose utilization and fitness through mutations in CSE2 (a subunit of RNA Pol II mediator), IRA1 (the Inhibitory Regulator of the RAS-cAMP pathway), MTH1 (involved in glucose-regulated gene expression), UBR1 (an E3 ubiquitin ligase), and ACE2 (involved in full septation of budding)^20^ (**Supplementary Fig. 3**). These mutations collectively led to increased local cell densities, which enhanced cell growth via extracellular sucrose cleavage^20^. Additionally, because our induction system (pBAD) is under the control of arabinose—a carbon source for both *E. coli* and *S. cerevisiae*—we knocked out arabinose utilization from the *E.coli* strain (PAS842, see Methods for details) and grew the *S. cerevisiae* strain (PAS844) in anaerobic conditions such that arabinose was not consumed^21^.

To verify whether the sucrose titers were sufficient to grow heterotrophs, we first grew *E. coli* and *S. cerevisiae* in the spent lithotrophic minimal media from 9 days of PAS837 growth with and without induction of the sucrose synthesis genes. The *E. coli* was inoculated at OD_600_ = 0.01 and *S. cerevisiae* at OD_600_ = 0.05 and grown anaerobically for 48 h at 30 °C while shaking. Both *E. coli* and *S. cerevisiae* cultures grow in spent media from induced PAS837 by two doublings but showed no growth in uninduced PAS837 (later supplemented with 0.3% arabinose during heterotroph growth) (**Supplementary Fig. 4**). This result validated that engineered *C. necator* could support the growth of these heterotrophs.

We focused on *E. coli* co-culture because *C. necator* is aerobic, thereby preventing the further use of our *S. cerevisiae* strain. Once *C. necator* PAS837 was stably growing in lithotrophic conditions (30:15:balance, H_2_:CO_2_:air), we back-diluted to OD_600_ = 0.5, allowed cells to grow for 72 h, at which time the *C. necator* cultures were both induced and co-inoculated with *E. coli* PAS842 and allowed to grow for an additional 7 days (**Fig. 2b,d,f**). The *E. coli* PAS842 in co-culture with induced *C. necator* PAS837 grew to 22x higher cfu/mL compared to WT *C. necator* and 2x higher compared to uninduced *C. necator* PAS837. Growth of *E. coli* in induced co-culture coincided with a sharp decrease of sucrose measured in the supernatant, from 60 mg/L to less than 10 mg/L (**Fig. 2b, d**). We did not observe sucrose in the supernatant of either WT or uninduced *C. necator* co-cultures. Overall, *C. necator* WT cfu/mL was higher compared to engineered strains and all growth declined over time (**Fig. 2f**).

Notably, the growth of *E. coli* in co-culture with *C. necator* exceeded expected growth based on monoculture sucrose production by *C. necator*. In order to determine *E. coli* sucrose requirements, we grew *E. coli* in increasing concentrations of sucrose and correlated resulting *E. coli* cfu/mL after 2 days (**Fig. 2c**). This method allowed us to compare *E. coli* growth in supernatant with growth in co-culture. *E. coli* growth in co-culture corresponded to 2.5-fold higher sucrose concentrations than those measured in *C. necator* monocultures (**Fig. 2c**). The growth patterns we observed are consistent with a model in which a symbiotic heterotroph acts as a thermodynamic sink for feedstock production^22^—a possible explanation for the increased growth and higher sucrose production. Co-culturing a heterotroph directly with a feedstock producer could therefore be a method to enhance both the heterotroph’s growth and its potential product yield.

Finally, we showed that *E. coli* can produce violacein in co-culture with sucrose producing *C. necator*. We introduced violacein pET21b-VioABCDE^23^ (PAS845) and β-carotene-expressing pSB1C3-CrtEBIY^24^ (PAS846) plasmids to our *E. coli* strain. For production we grew 100 mL of *C. necator* lithotrophically for 3 days before induction and addition of violacein- or β-carotene-producing *E. coli* at OD_600_ = 0.01. Gas was replenished every three days and the cultures were harvested 10 days after induction. After cultivation, the cells were harvested by centrifugation and products were extracted. Violacein and β-carotene were quantified by comparison to an analytical standard. The cells produced 80–250 µg/L violacein and 50 µg/L carotene from CO_2_ (**Fig. 2e**). *E. coli* grew by about 2 orders of magnitude while *C. necator* growth was reduced by one half of an order of magnitude. An important caveat in the reported titers is that due to H_2_ safety requirements the *C. necator* strains rarely exceeded densities above OD_600_ of 3. In sum, violacein and beta-carotene are produced from CO_2_ and H_2_ as sole sources of carbon and energy in this system.

### Polyhydroxyalkanoate (PHA) production

Extensive work has been done to understand and optimize *C. necator* PHA bioplastic production and has achieved yields up to 1.5 g/L/h in lithotrophic conditions^25^. The most commonly produced PHA, polyhydroxybutyrate (P(3HB), PHB), is a brittle thermoplastic with a narrow processing window and suboptimal material properties^8^. To extend the utility of *C. necator* for bioplastic production, we combined gas fermentation and genetic engineering to tailor more versatile PHAs from CO_2_ and H_2_.

To produce these tailored biopolyesters, we expressed thioesterases (TEs) with PHA synthases (phaCs) (**Fig. 1b**). As the last step in PHA synthesis, phaCs assemble the polymer based on their intrinsic substrate specificities and enzymatic rates. Introducing heterologous enzymes with variable specificities for different chain-length fatty acids, via TEs, into this process should thus generate bioplastics with tailored compositions. Further alterations to composition and production can be made by either removing or complementing the native PHA synthase (phaC*_Cn_*) or modulating β-oxidation.

PHA copolymers are generally derived through the introduction of the co-monomer precursors as feedstocks, of which saturated fatty acids are the most common. Rather than feeding these precursors, we introduced TEs to produce fatty acids to then be integrated into the copolymer. We selected two acyl carrier protein (ACP) TEs: plant TE *U. californica* FatB2 a 12:0 acyl-ACP TE^26,27^ and our engineered chimera of *C. palustris* FatB1(aa 1-218) and FatB2 (aa 219-316)—Chimera 4 (chim4)^28^. Chim4 couples the specificity of *Cp*FatB1 as a C8:0 acyl-ACP TE with the high activity of *Cp*FatB2, which is natively a predominantly C14:0 TE. Previous work established that when overexpressed in *E. coli*, *Cp*FatB1 produced 84 µg/mL of C8:0 (78% total FFA produced) and *Cp*FatB2 372 µg/mL C14:0 (75% total FFA). The engineered chim4 produced 394 µg/mL of C8:0 (90% FFA)^28^. By combining these TEs with phaC polymerizing enzymes, we were able to produce PHA copolymers directly from CO_2_ that were equivalent to those made from heterotrophic conditions. The three phaCs that we used were: *P. aeruginosa* phaC1 and phaC2 and *Pseudomonas spp 61-3* phaC1*_Ps_*, which were co-expressed with the ACP TEs to generate the medium-chain length (mcl) PHAs. As reported in the literature, when phaC1*_Pa_* is overexpressed in PHA-producing *E. coli* and fed C12 fatty acid, the PHA copolymer was composed of approximately 45% C12, 50% C10, 10% C8 molar proportion^29^. In phaC2*_Pa_*-overexpressing *E. coli* when fed C12 fatty acid a polymer composed of 35% C12, 55% C10, 10% C8 hydroxy acids was produced. When this *E. coli* strain was fed C8 fatty acid, the polymer was instead composed of 15% C10, 40% C8, 35% C6, 10% C5 hydroxy acids^30,31^. When fed C8 fatty acid, *P. putida* GPp104 expressing the PHA synthesis pathway with *Pseudomonas spp* 61-3 phaC1_Ps_ produced 2% C10, 77% C8, 16% C6, 3% C4 hydroxy acids^32^.

During lithotrophic growth, *C. necator* strains produced a variety of expected copolymers based on the combination of genetic background, the genes of the expression plasmid, and pharmacological manipulation (e.g., β-oxidation inhibition^33^). The PHA was purified through established NaClO^−^-based protocols^34^ and underwent subsequent methanolysis before GCMS analysis of medium chain-length hydroxy acids (mcl-3HA) at m/z =103 (**Supplementary Fig. 1b**). PHA titers based on percent dry cell weight (%DCW) are reported in **Supplementary Fig. 6**., all data are representative of n = 3 (all subsequent percentages are reported as the mean from n = 3 unless otherwise noted). *C. necator* with the vector control ΔphaC*_Cn_* (PAS827) failed to produce detectable PHAs, whereas WT cells (PAS826) produced 100% 3HB; with no effect from acrylic acid in either condition (**Fig. 3a**). In the native phaC*_Cn_* background, this enzyme pair (PAS828) produced very little mcl-3HA: μ = 4.4% of total (**Fig. 3b**, first panel, **Supplementary Fig. 7a**). In the ΔphaC*_Cn_* background (PAS829), this plasmid produced μ = 79.7% mcl-3HA, of which 64.5% was 3HO (**Fig. 3b**, second panel). Acrylic acid did not contribute significantly to the PHA composition; when added to PAS828, the proportion of 3HO and 3HD decreased to undetectable levels (**Fig. 3b**, third panel). PAS829 in the acrylic acid condition, produced a copolymer that was slightly enriched for longer-chain fatty acids compared to no-acrylic acid condition (**Fig. 3b**, fourth panel). Overall, while there were striking differences between the native and ΔphaC strains, acrylic acid did not seem to affect the proportion of fatty acids in this strain.

**Figure 3:**
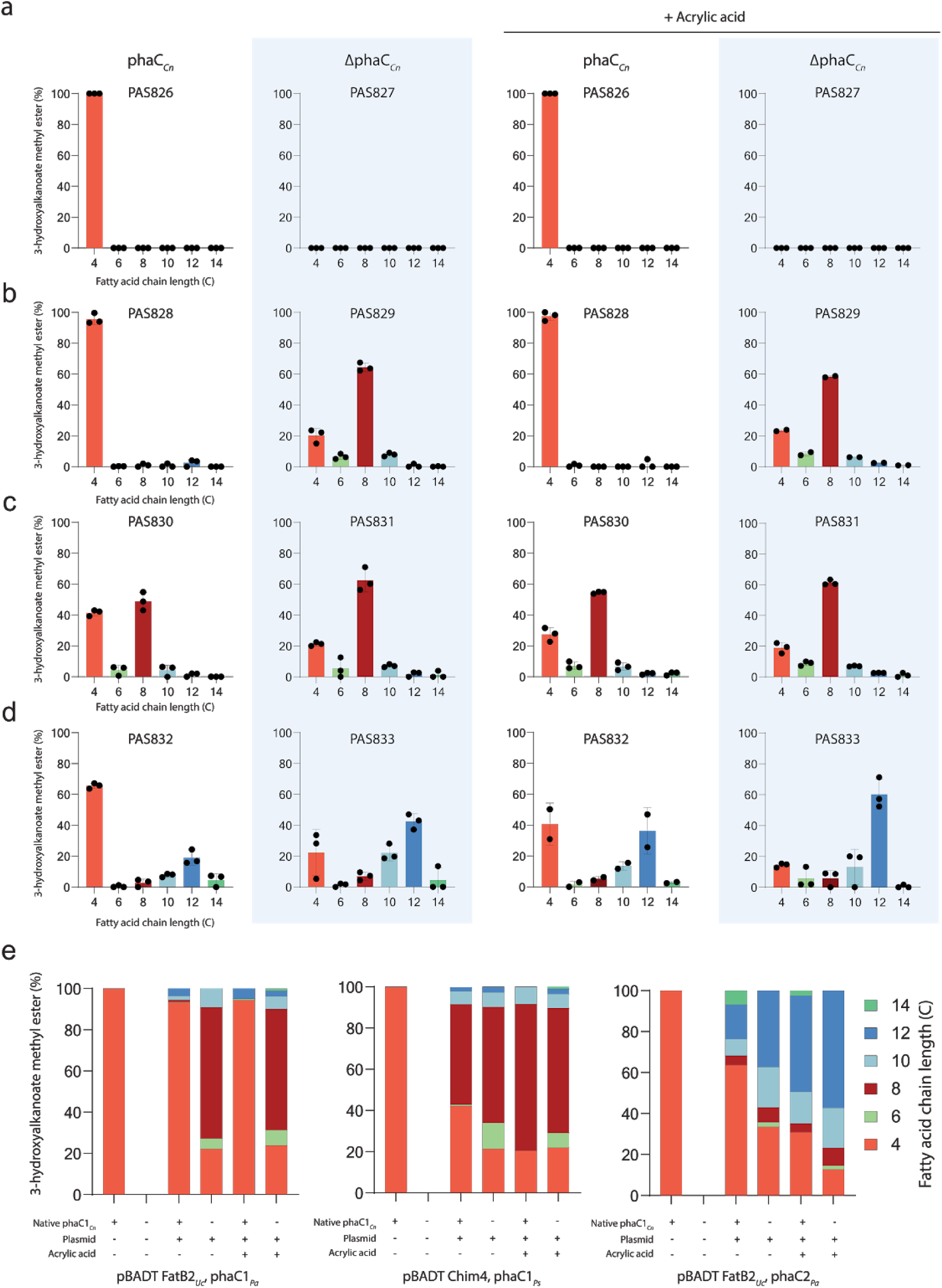
Lithotrophic production of tailored PHA. 3-hydroxyalkanoate mol/mol ratio from methanolyzed methyl esters of the purified PHA. Values are based on AUC for each peak at m/z=103 via GCMS (n=3 for each condition). (**a-d**) The first and third columns are strains with native phaC*_Cn_* background; the second and fourth columns are in the knock-out ΔphaC*_Cn_* background. The third and fourth columns are with the administration of 240 μg/mL acrylic acid with induction. (**a**) Panel one, WT phaC*_Cn_* produces 100% 3HB. Panel two, knock-out ΔphaC*_Cn_* does not produce detectable amounts of PHAs. Third and fourth panels, the composition of the polymer is not strongly affected by acrylic acid in either strain, all data set are n=3 unless otherwise noted. (**b**) PAS828 (phaC*_Cn_*, pBAD UcFatB2, phaC1*_Pa_*); PAS829 (ΔphaC*_Cn_*, pBAD UcFatB2, phaC1*_Pa_*), fourth panel (n=2) (**c**) PAS830 (phaC*_Cn_*, pBAD chim4, phaC1*_Ps_*) PAS831 (ΔphaC*_Cn_*, pBAD chim4, phaC1*_Ps_*). (**d**) PAS832 (phaC*_Cn_*, pBAD UcFatB2, phaC2*_Pa_*) PAS833 (ΔphaC*_Cn_*, pBAD UcFatB2, phaC2*_Pa_*), third panel (n=2). (**e**) Side-by-side comparison of representative copolymers in each condition to demonstrate the predictable trends of those conditions. Fatty acids represented by: C4, bright red; C6, light green; C8, dark red; C10, light blue; C12, royal blue; C14, teal.

The second group of experiments used *Pseudomonas spp 61-3* phaC1*_Ps_* and our engineered chim4 TE enzyme with high activity and selectivity for octanoate production (**Supplementary Fig. 5**)^28^. This plasmid in the native phaC background (PAS830) produced μ = 58.5% mcl-3HA, of which μ = 48.9% was 3HO (**Fig. 3c**, first panel, **Supplementary Fig. 7b**). Surprisingly, in the ΔphaC*_Cn_* background (PAS831) the composition of mcl-3HAs compared to the native background (PAS830) was not as striking as PAS828 to PAS829; with μ = 78.7% being mcl-3HA with μ = 62.6% as 3HO (**Fig. 3c**, second panel). We observed the strongest effect of acrylic acid on a strain with the native phaC, with the accumulation of longer chain fatty acids: μ = 72.6% mcl-3HA, of which μ = 54.6% was 3HO (**Fig. 3c**, third panel). After the addition of acrylic acid, we saw a minor enrichment of mcl-3HAs in PAS831: μ = 61.4% 3HO, μ = 7.1% 3HD, μ = 2.6% 3HDD, μ = 1.2% 3HTD (n = 2) (**Fig. 3c**, fourth panel).

In a third set of experiments, we explored the tunability of different combinations of TEs and phaCs by exchanging the phaC1*_Pa_* with its paralog phaC2*_Pa_* while maintaining the *Uc*FatB2. The plasmid containing TE *Uc*FatB2 and phaC2*_Pa_* in the native phaC*_Cn_* background (PAS832) mcl-3HAs comprised approximately μ = 34.5% of the polymer, of which μ = 18.9% of 3-hydroxydodecanoate (3HDD) (**Fig. 3d**, first panel, **Supplementary Fig. 7c**).

In strains lacking the native phaC*_Cn_* and expressing the above plasmid (PAS833), a greater accumulation of mcl-3HA was observed; μ = 77.6% was mcl-3HA, of which μ = 42.5% was 3HDD (**Fig. 3d**, second panel). When acrylic acid, an inhibitor of 3-ketoacyl CoA thiolase in β-oxidation, was added to PAS832, mcl-3HAs accumulated to a similar extent as seen in the ΔphaC strain (PAS833) without acrylic acid, μ = 59.4 with μ = 36.3% 3HDD (n = 3) (**Fig. 3d**, third panel). Likewise, in PAS833, a marked increase in the proportion of 3HDD predominated; 60.3% 3HDD (**Fig. 3d**, fourth panel).

Together, these experiments indicate we can modify the composition of the PHA polymer in many predictable ways (**Fig. 3e**).

### Lipochitooligosaccharide production

We engineered *C. necator* to convert CO_2_ into lipochitooligosaccharides (LCOs), a plant growth enhancer^35–37^. We used a pBAD-based plasmid containing NodC protein, an N-acetylglucosaminyltransferase that builds the backbone, NodB, a deacetylase that acts on the non-reducing end, and NodA, an acetyltransferase that attaches a fatty acid (**Fig. 1c**). This pathway produced a basic LCO molecule composed of a 5 (V) acylated chitin backbone and oleic acid at the non-reducing terminal N-acetylglucosamine monomer, which, following naming convention, we called Nod Cn-V (C_18:1_).

We initially detected the Nod Cn-V (C_18:1_) from our engineered strain by HPLC and used purified *B. japonicum* Nod Bj-V (C_18:1_ MeFuc) as a reference. Consistent with the literature, we observed two peaks at the expected retention times (**Supplementary Fig. 8**). In lithotrophic conditions, the engineered *C. necator* produced 1.37 ± 0.44 SE mg/L over 72 h (**Fig. 4a**). LC-MS analysis confirmed the identity of Nod Cn-V (C_18:1_). While all fragmentation peaks were consistent with Nod Bj-V (C_18:1_ MeFuc) (m/z = 1035, 831, 629, and 426), the base peak at m/z = 1256 indicates the lack of the fucose moiety^35,38^ (**Fig. 4b** and **Supplementary Fig. 9**).

**Figure 4.**
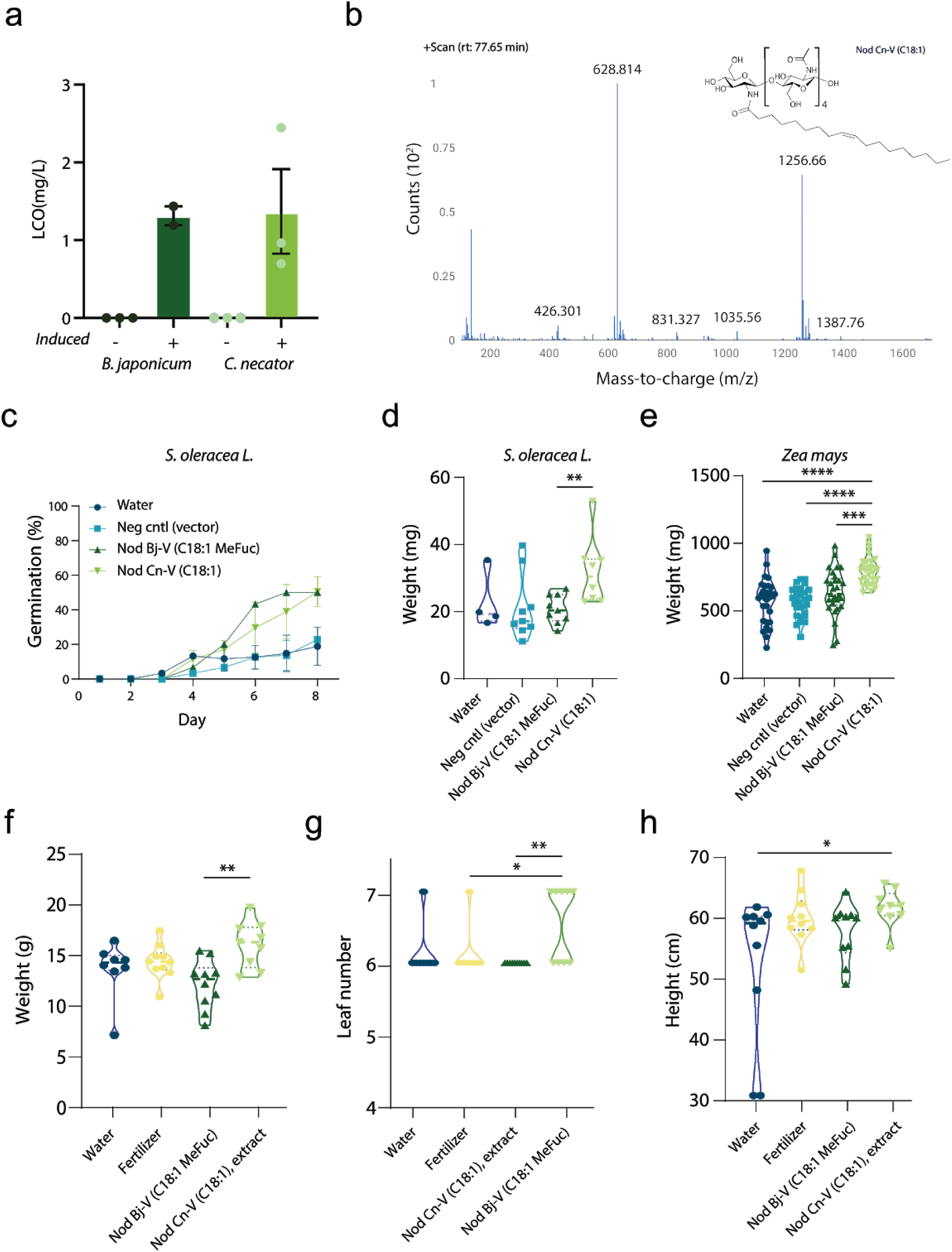
Lithotrophic production of Nod Cn-V (C_18:1_) in *C. necator*. (**a**) Yields of WT *B. japonicum* strain 6 (dark green) and LCO-producing *C. necator* (light green) +/- inducer (genistein and arabinose, respectively) (n=3). (**b**) LCMS mass spectrum of eluted peak at 77.65 min containing Nod Cn-V (C_18:1_). (**c**) Germination rates in seeds in response to LCO application for spinach. Seeds were treated with: water (dark blue circles), a vector control (blue squares), a standard LCO (Nod Bj-V (C_18:1_ MeFuc)) control from *B. japonicum* (dark green up-triangles) and the extract from *C. necator* vector control (bright green down-triangles) (n=10 per experiment, which were done in triplicate). (**d**) Spinach germination weight increased in Nod Cn-V (C_18:1_) compared to Bj-V (C_18:1_ MeFuc) (p=0.0072), not all seeds germinated which is reflected in the different number of samples in each group: water (n=4); negative control (n=9); Nod Bj-V (C_18:1_ MeFuc) (n=9); Nod Cn-V (C_18:1_) (n=8). Two-sided Mann-Whitney test was used. (**e**) Corn germination weight increased significantly in Nod Cn-V (C_18:1_) compared to all conditions: water (p<0.0001), vector control (p<0.0001), and Nod Bj-V (C_18:1_ MeFuc) (p=0.0003). Water (n=30); negative control (n=29); Nod Bj-V (C_18:1_ MeFuc) (n=31); Nod Cn-V (C_18:1_) (n=28). Two-sided Mann-Whitney test was used. (**f**) Growth characteristics of greenhouse corn. Nod Cn-V (C_18:1_) extract samples were derived from 1-butanol extraction. Nod Cn-V (C_18:1_) increased corn wet weight compared to Nod Bj-V (C_18:1_ MeFuc) (p=0.0023) and Nod Cn-V (C_18:1_), (**g**) Nod Cn-V (C_18:1_) significantly increased leaf number compared to fertilizer (p=0.036), Nod Bj-V (C_18:1_ MeFuc) (p=0.007). (**h**) Nod Cn-V (C_18:1_) significantly increased the corn height as compared to water (p=0.04). Not all plants grew, which is reflected in the number of samples in each group: water (n=8); fertilizer (n=10); Nod Bj-V (C_18:1_ MeFuc) (n=10); Nod Cn-V (C_18:1_) (n=9). Violin plots depict median (dashed line) and quartiles (dotted line). Multiple comparisons one-way ANOVA were used for (4e-h). Asterisks indicate significance: * indicates <0.05, ** indicates <0.01 and *** indicates <0.001.

Following quantification and spectrometric analysis, purified Nod Cn-V (C_18:1_) was applied to the seeds of different plant species (**Supplementary Fig. 1c**). We used spinach, corn, and soybean for their fast growth rates and to remain consistent with the literature. Because plant responses to LCOs are variable, we tested a range of concentrations for each species. After a germination time of 8 days, the engineered Nod Cn-V (C_18:1_) increased the percent germination for spinach seeds by 125%, with an average of 60% seeds germinated compared to 40% for vector control and 27% for water as a control, respectively (**Fig. 4c** and **Supplementary Fig. 10**). Nod Cn-V (C_18:1_) had no effect on corn or soybean germination (data not shown). In addition to percent germination, Nod Cn-V (C_18:1_) increased growth parameters in spinach. Nod Cn-V (C_18:1_) significantly increased the overall spinach sprout weight compared to the vector control (p < 0.05), with a 41% increase compared with Nod Bj-V (C_18:1_ MeFuc) (**Fig. 4c**).

No significant difference was found for spinach length (data not shown). Germinated corn sprout weight with Nod Cn-V (C_18:1_) was significantly increased by 42% as compared with water (p<0.0001), vector control (p<0.0001), and Nod Bj-V (C_18:1_ MeFuc) (p=0.0003) (**Fig. 4e**). Because all of the corn germinated and sprouted during the experiment while the spinach did not, we have more samples for corn. Nod Cn-V (C_18:1_) significantly increased corn shoot weight by 55% compared to water (p < 0.05) and shoot length by 25% compared to water (p = 0.002) (**Supplementary Fig. 11**). No differences were found for corn root weight or length (data not shown). No significant differences were found in soybean (data not shown).

In addition to germination experiments, we observed that the application of Cn-V (C_18:1_) increased corn yield in greenhouse crop growth experiments (**Supplementary Fig. 12**). Corn wet weight increased significantly with Nod Cn-V (C_18:1_) by 18% when compared with water (p=0.0023) (**Fig. 4f**). Nod Cn-V (C_18:1_) also significantly increased corn leaf number when compared to fertilizer (p=0.036), Nod Bj-V (C_18:1_ MeFuc) (p=0.007) (**Fig. 4g**). Thirdly, Nod Cn-V (C_18:1_) significantly increased corn height compared to water (p=0.04) and was equivalent to the fertilizer condition (**Fig. 4h**). Nod Cn-V (C_18:1_) shows that it can lead to increased germination and growth in spinach, as well as increased growth of corn shoots and whole plant, generally outperforming the leguminous Nod Bj-V (C_18:1_ MeFuc) control. This suggests that our engineered LCO is a more generalizable plant-growth enhancer than the standard from *Bradyrhizobium spp*.

## Discussion

Here, we seek to further develop engineered lithoautotrophs to minimize agricultural land use and counteract pollution caused by petrochemical and agricultural industries. We established the plausibility of our approach by producing feedstocks for bioproduction, biodegradable bioplastics to offset petrochemical plastics production and pollution and plant-growth enhancers to promote increased food crop yields and efficiency while offsetting the use of synthetic fertilizers. Decoupling bioproduction from plant-based feedstocks will reduce the competition for crop land use, promote commercialization by lowering costs, and support the existing infrastructure of industrial heterotrophs^2^.

We have demonstrated that *C. necator* is a versatile strain to use in lithotrophic growth. We engineered the bacteria to make sucrose and support growth of heterotrophs, expanded the scope of gas-based PHAs, and improved the growth of plants through production of general purpose LCOs.

While co-culture systems have been developed with cyanobacteria with *E. coli* and *S. cerevisiae*^19^ as well as the acetogen *M. thermoacetica* with *Y. lipolytica*^22^, our system combines the higher energy feedstock, sucrose, with non-photosynthetic gas fermentation in a single reactor. Our results show that engineered heterotrophs can produce complex products in co-culture with *C. necator* without deleterious growth effects (see Supplemental discussion).

We produced a spectrum of de novo PHA copolymers from CO_2_ and H_2_. We used three primary levers with which to tailor the composition: overexpression of TEs to phaCs, the presence or absence of the endogenous phaC1*_Cn_*, and the addition of acrylic acid—a β-oxidation inhibitor. We demonstrated the feasibility with standard medium chain-length fatty acids thereby showing the potential to genetically control the composition of the copolymers such that specific and industrially relevant PHAs can be produced lithotrophically.

LCOs—unlike synthetic fertilizers that are volatile and require high concentrations—act at nanomolar to micromolar concentrations. While there have been some attempts to engineer *B. japonicum*, it remains challenging and still in the early stages^39^. We have demonstrated that *C. necator* is a viable chassis for LCO production. Previous work has enhanced LCO production in native rhizobia strains^35,38^, but there have been several unsuccessful attempts to produce LCOs through chemical synthesis^40^ and engineered *E. coli*^41^. A recent study produced LCOs using a combination of chemical and biological methods, but this approach is unlikely to be scalable or sustainable^42^. Our approach, inspired by mycelial LCOs with which over 60% of all plants are able to form arbuscular mycorrhiza associations^43^ and can promote growth^36^, sought to produce a general purpose LCO to apply to a variety of non-legume plants. Our design includes only the genes that are necessary and sufficient for building a functional basic Nod LCO. While we have demonstrated generality, a potential strength of this approach is that through genetic engineering the LCOs can also be tailored to a plant of interest for more effective growth enhancement. Use of *C. necator* allows for a more tractable genetic chassis that can grow to higher cell densities and faster rates than for example *Bradyrhizobium spp*.

Gas fermentation is progressing from a promising idea into an implementable industrial platform. Advances in fermenter construction^44–48^ have enabled the operation of large industrial bioproduction plants on site with syngas (CO, CO, H) point sources^4^ at minimized costs. In our proposed system, projected CO_2_ costs would be negligible for gas fermentation through the use of concentrated, mostly pure CO_2_ point sources such as breweries and ethanol-production plants^4^. Pure H_2_—produced by steam reforming or water electrolysis—would be the primary driver of feedstock cost. In addition to cost, it is important to consider the thermodynamic efficiencies of each process. While anaerobic methanogens and acetogens are extremely efficient at reducing CO/CO_2_, that efficiency sharply declines once the product becomes more complex than ethanol or acetate. *C. necator* produces 8-fold more ATP per H_2_ than methanogen or acetogens, and 4-fold more biomass per CO_2_ via the Calvin-Benson-Bassham Cycle^44^. Comparing H_2_ to plant-derived carbohydrates as feedstocks, the energy-to-feedstock efficiency is approximately 0.1% for plants and 14% for solar H_2_^45,46^.

Given this difference, we see the potential for *C. necator* systems to expand bioproduction beyond sugar-based feedstocks. Despite their prevalence, plant-derived sugars have hidden environmental costs to the planet. Their low price is born, in part, from a highly developed industrial agriculture—which has significant GHG emissions^47,48^. While microbes hold promise for sustainable intensification of agriculture^49^, the reliance of bioproduction on plants limits the global green economy—as issues of food security increase, the tradeoff between food and bioproducts will become increasingly difficult to make^50^. Using a CO_2_ -based gas fermentation, we have expanded the scope of a lithotrophic microbial chassis.

### Sustainability considerations

*C. necator* is one of the most effective microbes in converting H_2_ into biomass. The solar-to-biomass efficiency for terrestrial plant photosynthesis is 1% with only 10-20% conversion efficiency of CO_2_ into sucrose itself (e.g, sugar beets, sugarcane), compared to 3-5% and 80% for cyanobacteria, respectively, and 18% and 11% for *C. necator*, respectively (**Fig. 5a**). Lithoautotrophic production is non-photosynthetic and is not limited by the thermodynamic and technological limitations of plant and cyanobacteria-based bioproduction systems^19,56^. Here, based on general assumptions in the literature and using solar radiation as the common energy source, we estimate that for 90 g/L biomass—typical yield for *C. necator* grown in a (CO_2_/O_2_/H_2_) gas fermentation system—with our reported production efficiency of 11.3%, we can produce 510 tons of sucrose per hectare (ha) of photovoltaics per year (details in the Supplemental Methods). These estimates, based on equivalent land use, indicate that our approach is 35-fold more productive than plants and 13-fold more productive than cyanobacteria (**Fig. 5a**).

**Figure 5.**
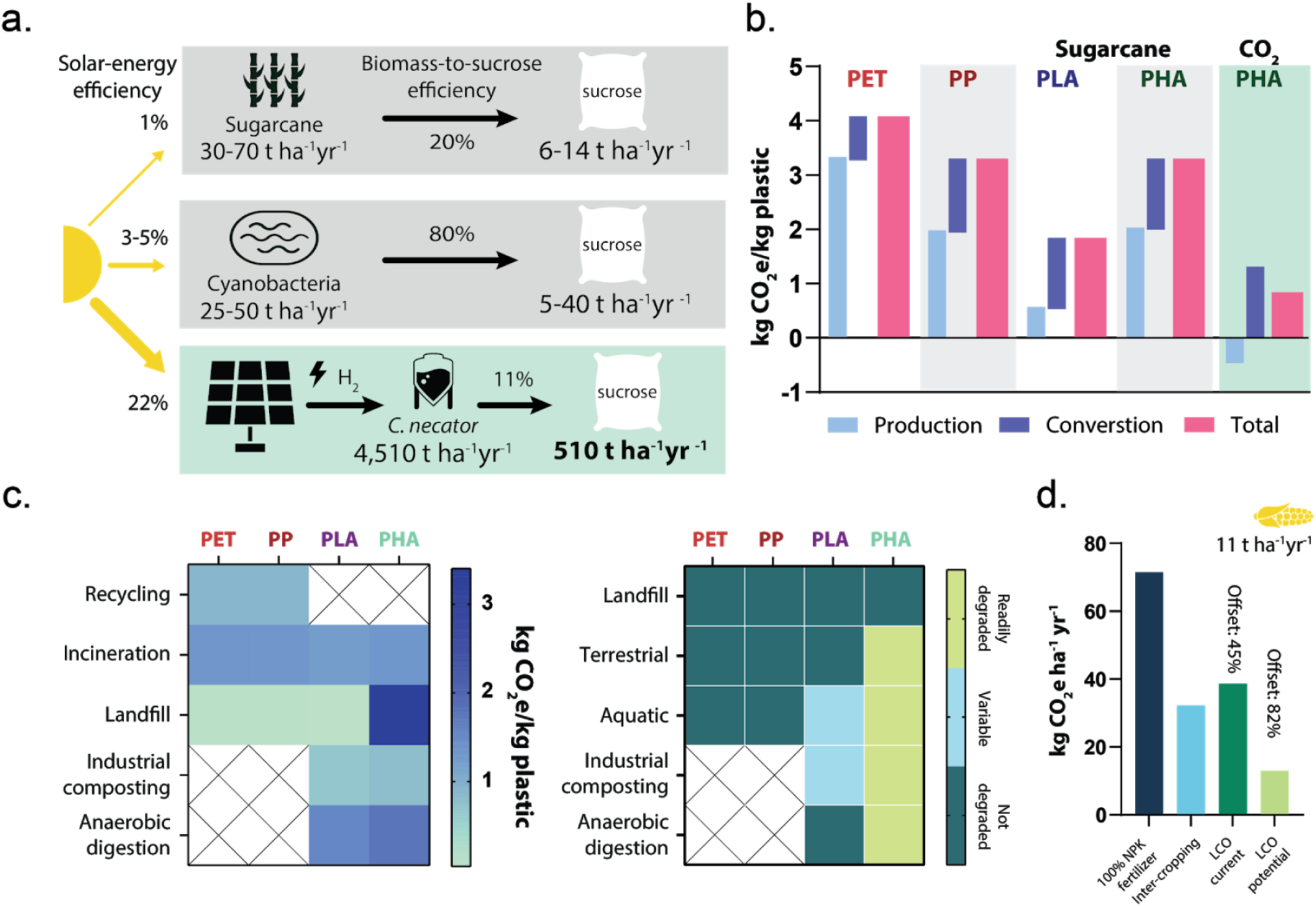
Sustainability comparisons of existing strategies. (**a**) Comparison between sugarcane, cyanobacteria, and *C. necator*. Solar-to-biomass conversion efficiency of plants is approx. 1% annually; cyanobacteria is 3% in open ponds and 5-7% in photobioreactors. Photovoltaics (PVs) with an average solar-to-energy conversion of 22% can generate H_2_ at 14% efficiency. 30-70 ton ha^−1^yr^−1^ of sugarcane biomass with 20% biomass-to-sucrose efficiency converts to 6-14 ton ha^−1^yr^−1^ sucrose. For cyanobacteria, with an 80% biomass-to-sucrose efficiency, 5-40 ton ha^−1^yr^−1^ of sucrose. For *C. necator*, using PV area as the land use equivalent and using our biomass-to-sucrose efficiency of 11%, 4,510 ton ha^−1^yr^−1^ of biomass can produce 510 ton ha^−1^yr^−1^ of sucrose. (**b**) GHG emissions for main classes of plastics (PET, polyethylene terephthalate; PP, polypropylene; PLA, polylactic acid; PHA, polyhydroxyalkanoates) with current energy mix^51^. Based on our conversion efficiency of PHA 50% DCW and the CO_2_ drawdown rate of 0.61-0.65 g of biomass per g CO_2_. (**c**) First panel; the carbon footprint of the end-of-life of plastics^51,52^. Second panel; the relative biodegradability of main classes of plastics. Petrochemical plastics are not processed in industrial composter or anaerobic digestion. Depending on conditions of the composters/digesters, PLA will not degrade. (**d**) Fertilizer (NPK, nitrogen-phosphorus-potassium) offset from LCO supplementation. Based on average corn production of 11.1 ton ha^−1^yr^−1^ and 40% yield increase conferred by fertilizer, which served at 100% of possible growth increase. Intercropping uses different crops to increase soil quality. CO_2_e values are based only on NPK offset to produce equivalent yields^53–55^. LCO current values are based on field study yields. LCO optimized values are based on optimized greenhouse growth conditions. See additional methods and references in Supplementary Information.

When considering plausible sustainable alternatives to petrochemical plastics, several factors must be considered: material properties, cost of production, carbon footprint of production, and end-of-life. PHAs, unlike commodity bioplastics such as PLA, are polymerized biologically. Manipulation of PHA composition and, consequently, material properties can be made through genetic engineering. The importance of making industrially relevant polymers from gaseous feedstocks is to reduce costs and the carbon footprint of production. PLA, for example, is primarily derived from corn—which while it sequesters CO_2_—the carbon footprint of the industrial agricultural supply chain and polymerization negates much of the benefits of CO_2_ drawdown. Similarly, PHAs produced from carbohydrate feedstocks—from the perspective of carbon emissions—are likely to be less sustainable than their petrochemical analogs (**Fig. 5b**).

End-of-life concerns are relevant to consolidating the GHG emissions of waste processing as well as the accumulation of pollution in the environment. Balancing the different environmental impacts of plastics will be crucial to choosing the most sustainable alternatives. Given the highly compacted and anaerobic environment of landfills, biodegradation occurs slowly^57,58^. In the case of PHAs, anaerobic methanogens degrade the polymers into methane, which could ideally be captured and purified for use as a fuel source (**Fig. 5c**, first panel). While dependent on a variety of factors including composition and physical dimensions, petrochemical plastics and PLA that find their way into aquatic and terrestrial environments persist on long, and in the case of nanoplastics, potentially indefinite timescales^59^ (**Fig. 5c**, second panel). As of 2010, 4.8-12.7 Mt of plastic persist in the oceans, and by 2025 estimates predict 90-250 Mt^60^. Plastic pollution permeates our planet^61^ and is a major threat to global biodiversity^62^. Constructing a mitigation strategy that addresses GHG emissions and pollution will be vital to a sustainable economy.

We see gas fermentation as a mechanism to enhance agricultural productivity. In the context of climate change—and the ensuing reduction of arable land and crop yields—in combination with a push towards greener industries, we must consider the bioproduct-vs-fuel dilemma for commodity bioproduction^63^. Synthetic fertilizers currently support the production of food for half of the world’s population food requirements^64^ but also use 2% of the global energy and produces 3% of the global GHG emissions^65^. In addition to GHG emissions from its synthesis, nitrogen once in the environment leads to long-term ecological damage^66,67^. In an effort to offset synthetic fertilizer production with bioproduction to minimize competition for land use, we seek to improve the efficiency with which agriculture can respond to increasing demand for plant-derived food, fuel, and textiles. A potential solution to more efficient and sustainable fertilizer use is the development of plant growth enhancers like LCOs (**Fig. 5d**).

Gaseous feedstocks like CO_2_ and H_2_ provide inexpensive and abundant sources of energy for industrial bioproduction. Our efforts seek to promote the use of gas fermentation to both offset sources of GHG as well as not compete with food for arable land.

## Methods

### Strain construction

All plasmid construction used Gibson Assembly in *E. coli* DH5α. Expression vector pBadT (JBEI) was conjugated into *C. necator* (ATCC 17699) from the donor strain MFDpir (generously provided by George Church’s lab). All genes, except the nodABC gene cluster, which was amplified from *B. japonicum* USDA 6^68^, were codon optimized and synthesized by SGI. *C. necator* knock-outs were constructed using integration vector pT18mobsacB via the conjugation methods described above and sucrose counterselection. *E. coli* W ΔcscR strain (generously provided by the Claudia Vicker’s laboratory^51^) and was transduced with the ΔaraC from the Keio collection, which conferred kanamycin resistance and removed the arabinose utilization operon through genetic linkage. *S. cerevisiae* W303^clump^ (generously provided by Andrew Murray’s laboratory^6^). Synthesized genes: *Umbellularia californica* FatB2 (12:0-ACP thioesterase); *Pseudomonas aeruginosa* phaC1; *Pseudomonas aeruginosa* phaC2; *Anabaena cylindrica* PCC 7122 SPS (sucrose phosphate synthase) and SPP (sucrose phosphate phosphatase); *Synechocystis sp.* PCC 6803 SPS (sucrose phosphate synthase) and SPP (sucrose phosphate phosphatase); *Escherichia coli* scrY (sucrose porin).

### Cell culture

Bacterial strains and growth protocols. Growth protocols follow the procedures in literature^13^. *E. coli* DH5α and MFDpir were grown in LB and LB with 300 µM diaminopimelate (DAP), respectively at 37°C. *C. necator* was grown at 30°C, for pre-culture in rich media (17.5 g/L nutrient broth, 7.5 g/L yeast extract, 5 g/L (NH_4_)_2_SO_4_). For lithotrophic growth, *C. necator* was grown in minimal medium was 3.5 g/L Na_2_HPO_4_, 1.5 g/L KH_2_PO_4_, 1.0 g/L (NH_4_)_2_SO_4_, 80 mg/L MgSO_4_·7H_2_O, 1 mg/L CaSO_4_·2H_2_O, 0.56 mg/L NiSO_4_·7H_2_O, 0.4 mg/L ferric citrate, and 200 mg/L NaHCO_3_. For nitrogen-limited growth to produce PHA, the (NH_4_)_2_SO_4_ concentration was reduced to 0.3 g/L. All solutions were filter-sterilized prior to use except the ferric citrate component, which was added after the filter sterilization step. Media was supplemented with 300 µg/mL kanamycin (PHA and LCO) or 25 µg/mL chloramphenicol (sucrose co-culture). The cultures were placed in a Vacu-Quick jar filled with H_2_ (8 inHg) and CO_2_ (2 inHg) with air as balance. Cultures were magnetically stirred and jars were refilled everyday with fresh gas mixture. To transfer the *C. necator* from heterotrophic to lithotrophic growth, an overnight culture grown in rich media was pelleted and washed twice with PBS, seeded in minimal media at OD_600_ = 0.2 and allowed to grow for 5 days until OD_600_ = 2. Cultures were then transferred into fresh minimal media and seeded at OD_600_ = 0.2. *Bradyrhizobium japonicum* strain 6 was cultured in a medium with 2.6 g/L HEPES, 1 g/L yeast extract, 0.5 g/L gluconic acid, 0.5 g/L mannitol, 0.22 g/L KH_2_PO_4_, 0.25 g/L Na_2_SO_4_, 0.3 g/L NH_4_Cl, 0.0112 g/L FeCl_3_ .6H_2_O, 0.017 g/L CuCl_2_ ·2H_2_O, 0.18 g/L MgSO_4_·7H_2_O, NaMoO_4_·7H_2_O, 0.0021 g/L NiCl_2_·6H2O, 0.01 g/L CaCl_2_·2H_2_O. Grown at a 30°C under continuous shaking at 200 rpm.

### Sucrose assay

75 mL of sucrose-producing *C. necator* cultures were grown lithotrophically as described above in 1 g/L (NH_4_)_2_SO_4_. Growth in co-cultures was monitored every 48 hours: OD_600_ was measured using the Ultrospec 10 Cell density meter (Amersham Biosciences) and cell numbers were assayed by plating dilution series on rich media to count colony forming units (CFU). Cell numbers of W303 Clump were derived by counting CFUs, and numbers were adjusted for the ∼6.6 cells/ clump as previously reported^6^. At each timepoint, cultures were pelleted and sucrose was measured from the supernatant and lysed pellet fractions using sucrose/D-glucose assay kits (Megazyme). Biological replicates were performed in triplicate.

### Heterotrophic cross-feeding and co-culture with *C. necator*

Back-diluted *C. necator* from lithotrophic growth at OD_600_ = 0.5 into lithotrophic growth and let grow for 2 days. Induced with 0.3% arabinose and added pre-condition *E. coli* at OD_600_ = 0.01. Took samples and plated selectively for cfu/mL counts every other day. Biological replicates were performed in triplicate.

### Quantification of *E. coli* products

Violacein and β-carotene were extracted in 100% ethanol and quantified via absorption and standard curves. 100 mL or 75 mL of co-culture was harvested 7 days after induction, pelleted and re-suspended in 1 mL of 100% ethanol. The suspension was centrifuged and the supernatant containing the product was separated. The absorbance at 580 nm (violacein) and 440 nm (β-carotene) was measured and compared to respective standard curves with violacein (Millipore Sigma) and β-carotene (Millipore Sigma) for quantification.

### Growth and induction for thioesterase

For thioesterase experiments, cells were grown in minimal media (described above) with 5% fructose. Once the cells reached mid-log, they were induced with 0.5% arabinose along with 800 µL 1-octanol as the organic phase. Aqueous and organic phases were harvested after 24, 72, and 120 h for GC-MS analysis.

### Fatty acid identification and quantification using GC-MS

FFAs dissolved in the aqueous culture phase were harvested by acidifying 400 µL aqueous culture with 50 µl 10% (wt/vol) NaCl and 50 µL glacial acetic acid and extracting the FFAs into 200 µL ethyl acetate. 100 µL of the ethyl acetate phase was then esterified in 900 µL of a 30:1 mixture of ethyl alcohol (EtOH) and 37% (vol/vol) HCl by incubating at 55°C for 1 h, and the ethyl esters were extracted into 500 µL hexanes for GCMS analysis. FFAs dissolved in the 1-octanol culture phase were esterified by acidifying 100 µL 1-octanol culture with 10 µL 37% (vol/vol) HCl then incubating at 55°C for 1 h, and the octyl esters were extracted into 500 µL hexanes for GCMS analysis. Fatty esters were analyzed on an Agilent GCMS 5975/7890 (Agilent Technologies) using an DB-35MS column. Samples were heated on a gradient from 40 to 250°C at 5°C/min. FFA chain lengths were identified by GC retention times and the mass spectra of the octyl esters at m/z = 112 and quantified using an internal standard (800 mg/L pentadecanoate added to the culture before the extraction procedure). Known concentrations of C6, C8, C10, and C12 fatty acids were used to generate a standard curve and to quantify the production of single fatty acid species.

### PHA extraction and analysis

After 48 h of growth in the second media exchange, 100 mL lithotrophic PHA-producing *C. necator* cultures were induced with 0.3% arabinose where indicated along with 240 µg/mL acrylic acid where indicated and grown for 5 days at 30°C. Cultures were harvest, pelleted, and lyophilized overnight. Freeze dried pellets were weighed for dry cell weight. PHA was purified with 0.2 mL/mg DCW of 13% NaClO^−^ for 4 h at 30°C, washed twice with dH2O, washed once with acetone, then dried at 25°C overnight. 1:1 methanol and HCl in dioxane to a final volume of 3 mL (1% pentadecanoate as internal standard), the tubes were sealed with crimp top and incubated in oil bath at 90°C for 20h. Tubes were placed on ice, once cooled, 2 mL of chloroform was added and vigorously vortexed, 3 mL of dH2O was added followed by extensive vortexing, the organic phase was separated by centrifugation (10 min, 4,000 x g), organic phase removed and stored at −20°C until GCMS analysis, biological replicates were performed in triplicate.

### GCMS PHA analysis

PHA samples were analyzed by Agilent GC-MS 5975/7890 (Agilent Technologies) on a DB-35MS column. Samples were heated on a gradient from 40 to 250°C at 5°C/min. The copolymer composition was determined with the mass spectra of the 3-hydroxy alkanoic acid methyl esters at m/z = 103 and NIST Mass Spectral Library.

### LCO production and isolation

Overnight cultures were back-diluted into 50 mL cultures, which were grown up to stationary phase and diluted into 200 mL cultures to an OD_600_ = 0.2. Cultures were grown until an OD_600_ = 1.0 (1-2 days) and induced by adding genistein (Sigma) to *B. japonicum* to a final concentration of 5 mM and 0.3% arabinose to *C. necator.* The cultures were incubated for an additional 76 hours. These cultures were then extracted with 0.4 vol of HPLC-grade 1-butanol by shaking vigorously for 10 min. The material was then centrifuged for 10 min at 4000 rpm. The upper butanol phase was separated and dried in a rotary evaporator under vacuum at 55°C (Yamato, New York, USA). The product was redissolved in 2 mL of 20% acetonitrile and analyzed by HPLC. A Vydac C18 reverse-phase column (Vydac, 5 mm, 250 x 4.6 mm) was used with a flow rate of 0.7 mL min -1. As a baseline, 30% acetonitrile was run through the system for at least 10 min prior to injection. The purification was done for 10 min in isocratic solvent A (water-acetonitrile, 70:30 [vol/vol]), followed by a linear gradient from solvent A to solvent B (water-acetonitrile, 50:50 [vol/vol]) for 25 min. The chromatographic peaks corresponding to the LCOs eluted between 24 and 28 min and were identified by comparison with an LCO standard, Nod Bj-V (C_18:1_ MeFuc), prepared from *Bradyrhizobium japonicum* strain USDA 523C (a generous gift from Professor D. Smith, McGill University). There were two LCO peaks that were detected using UV absorption at 206 and 214 nm.

### LCO analysis

The LCO sample structures were also confirmed by positive-ion QTOF mass spectrometric analysis using a tandem mass spectrometer and Collision-Induced Dissociation in combination with HPLC on a 10 μL aliquot. The method and mobile phases were the same as those used with the HPLC alone except the mobile phases had an addition of 1% formic acid to both LCMS-grade water and LCMS-grade acetonitrile. LCOs from the LCO standard, induced *B. japonicum*, and *C. necator* strains were all analyzed. The energy of the Cs^+^ ions was 25 keV, and the accelerating voltage of the instrument was set to 8 kV. Results were recorded using a scanning method from m/z 1415 to 1417 over 1 sec. The main compound for the standard and *B. japonicum* gives a peak at z = 1416.7 (M_1_H). The main compound for the *C. necator* strains give a peak at z = 1256.7 (M_1_H). Samples from induced and uninduced WT *C. necator* as well as uninduced vector control and engineered strains were all tested for LCOs using this method and none were found.

### Germination

Three species were tested; *Spinacia oleracea L.* (spinach) variety Regent, *Glycine max L.* (soybean), and *Zea mays L.* (corn) variety Trinity F1. Varieties were partially chosen for their relevance to agriculture in order to make this study as easily applied as possible. Seeds of all three species were surface-sterilized with 2% NaClO^−^ for 2 min, rinsed with sterile dH_2_O 10 times for spinach and 6 times for soybean and corn and then blotted dry. Each seed was soaked in the appropriate treatment solution for 30 min after being sterilized. A 9-cm diameter sterile petri dish was made with 1% noble agar and a 1 mm filter paper disk was placed on top of the agar. Five seeds were transferred onto the filter paper in six plates and five milliliters of dH_2_O (negative control), LCO standard (positive control), *B. japonicum* LCO extract (control), traditional N-based fertilizer (Miracle grow, 1 mg/mL), or solutions of purified extract from *C. necator* vector control strain or Nod factor producing strain at concentrations of 10^−5^, 10^−6^, 10^−7^, 10^−8^, 10^−9^ or 10^−10^ M were dispensed into each Petri dish. The Petri dishes containing spinach seeds were incubated at 21±2 **°**C and those containing corn and soybean were incubated at 25±2 **°**C, all in the dark. The number of sprouted seeds was obtained at least every 2^nd^ day for 9 days for spinach and 6 days for both soybean and corn. At the end of these time periods, each seed and sprout were weighed. The root and shoot systems were disconnected from the rest of the sprout when applicable and weighed separately. The root length and shoot length were measured for each seed that sprouted when appropriate by identifying the longest root or shoot and measuring this against a metric ruler. The number of roots was also counted for each seed when appropriate. Since overall growth rates were relatively low for spinach, only successful, sprouted seeds were included. Only successful sprouting and growth events with roots greater than 0.5 mm were considered. All experiments were repeated in triplicate and assays were performed blinded.

### Plant Yield/Growth

A greenhouse experiment was carried out using the same *Zea mays* (corn) variety Trinity F1 as in the germination experiment. Seeds were surface-sterilized with 2% NaClO^−^ for 2 min, rinsed with sterile dH_2_O 6 times, blotted dry, and each seed was soaked in treatment solution for 30 min. The growth tests were conducted in 6-inch diameter plastic pots. The pots were filled with growing media and seeds were planted just underneath the surface of the soil, with n=10 plants in each condition. Each pot received 25 mL of either dH_2_O, induced *B. japonicum* LCO, traditional N-based fertilizer (Miracle Gro, 1 mg/mL), solutions of purified extract from *C. necator* Nod factor producing strain at a concentration of 10^−7^ M or a solution from filtered cultures of the LCO producing strain at a concentration of 10^−7^ M. The culture solution was first lysed through a freeze thaw cycle and sonication for 20 min at high intensity and then filtered through a 0.22 mm filter. The pots were placed in Harvard University’s temperature controlled greenhouse at 25±2 **°**C during late June and July at atmospheric levels of CO_2_. Each treatment had ten replicates. After 3 days the pots were irrigated from above every 3 days. After two weeks, the number of leaves and the length of the longest shoot were measured with a metric ruler and above ground biomass was harvested. The wet weight was taken and the samples were lyophilized for 2 days until dry, the dry weight was measured. Experiments were performed blinded.

### Statistical Analyses

All statistical analyses were performed in Graphpad Prism. The multiple comparisons were done using one-way ANOVA with Tukey’s HSD post-hoc test to calculate differences between individual treatments were done using two-sided Mann-Whitney Test.

**Table 1.**

| Name | Species | Strain | Origin |
| --- | --- | --- | --- |
| PAS824 | *E. coli* | MG1655<br>RP4-2-Tc::[ΔMu1::aac(3)IV-ΔaphA-Δnic35-ΔMu2::zeo] ΔdapA::(erm-pir) ΔrecA | Geroge Church lab |
| PAS825 | *E. coli* | pBADT FatB1<sub>Cp</sub>, Fat2<sub>Cp</sub> (chim4) | [25] |
| PAS826 | *C. necator* | pBADT rfp | This study |
| PAS827 | *C. necator* | ΔphaC<sub>Cn</sub>, pBADT rfp | This study |
| PAS828 | *C. necator* | pBADT FatB2<sub>Uc</sub>, phaC1<sub>Pa</sub> | This study |
| PAS829 | *C. necator* | ΔphaC<sub>Cn</sub>, pBADT FatB2<sub>Uc</sub>, phaC1<sub>Pa</sub> | This study |
| PAS830 | *C. necator* | pBADT Chim4, phaC1<sub>Ps</sub> | This study |
| PAS831 | *C. necator* | ΔphaC<sub>Cn</sub>, pBADT Chim4, phaC1<sub>Ps</sub> | This study |
| PAS832 | *C. necator* | pBADT FatB2<sub>Uc</sub>, phaC2<sub>Pa</sub> | This study |
| PAS833 | *C. necator* | ΔphaC<sub>Cn</sub>, pBADT FatB2<sub>Uc</sub>, phaC2<sub>Pa</sub> | This study |
| PAS834 | *C. necator* | pBADT SPS<sub>Ac</sub>, SPP<sub>Ac</sub> | This study |
| PAS835 | *C. necator* | pBADT SPS<sub>Ac</sub>, SPP<sub>Ac</sub>, scrY<sub>Ec</sub> | This study |
| PAS836 | *C. necator* | pBADT SPS<sub>Se</sub>, SPP<sub>Se</sub> | This study |
| PAS837 | *C. necator* | pBADT SPS<sub>Se</sub>, SPP<sub>Se</sub>, scrY<sub>Ec</sub> | This study |
| PAS838 | *C. necator* | pBADT nodABC<sub>Bj</sub> | This study |
| PAS839 | *B. japonicum* | USDA 6 | [63] |
| PAS840 | *B. japonicum* | USDA 523C | Donald Smith lab |
| PAS841 | *E. coli* | WT; csc+ (ATCC 9637) | [29] |
| PAS842 | *E. coli* | WcscR::FRT | [29] |
| PAS843 | *S. cerevisiae* | S.cerevisiae W303 Ancestor strain | [25] |
| PAS844 | *S. cerevisiae* | S. cerevisiae W303Clump Recreated02 | [25] |
| PAS845 | *E. coli* | pET21b-VioABCDE | [17] |
| PAS846 | *E. coli* | pSB1C3-CrtEBIY | [18] |

## Acknowledgements

The authors thank Pavel Gorelik and Ofer Mazor at the HMS Research Instrumentation Core Facility for assistance in reactor design and fabrication. We also thank Michael Lewandowski at The Wyss Institute Materials Characterization Core for assistance with sample analysis. We also thank Matthew Henke for assistance with LC-MS analysis. S.N.N. thanks The Wyss Institute for Biologically Inspired Engineering and the Harvard Climate Change Solutions Fund for support. S.B. thanks NSF Graduate Student Fellowship for support. D.M.L. thanks Harvard’s David Rockefeller Center for Latin American Studies and the Elizabeth and William Patterson Fellowship for support. D.G.N. thanks DOE DE-SC0017619 for support. P.A.S. thanks the TomKat Foundation for support.

## Author contributions

S.N.N. and M.Z. wrote the manuscript, S.N.N., M.Z., S.B. and D.M.L., conceived of and performed experiments. D.T. assisted with experiments. S.N.N., M.Z., S.B., D.M.L., P.A.S. and D.G.N. conceived of experiments and provided edits to the manuscript.

