## Supplemental Information for "Valorization of CO_2_ through lithoautotrophic production of sustainable chemicals in *Cupriavidus necator*"

**Supplementary discussion** **2**
Comparison to cyanobacterial co-culture systems 2
Sucrose production calculations 3
Carbon footprint of plastics calculations 4
Carbon footprint of fertilizer calculations 5

**References** **6**

**Supplementary figures** **8**
Supplementary figure 1: Experimental setup 8
Supplementary figure 2: Comparison of cyanobacterial enzymes in *C. necator* 10
Supplementary figure 3: Sucrose growth titration 11
Supplementary figure 4: Heterotroph growth in *C. necator* supernatant 12
Supplementary figure 5: Octanoate production by *C.necator* 13
Supplementary figure 6: PHA content relative to dry cell weight (DCW) 14
Supplementary figure 7: 3HA ratios in tailored PHAs 15
Supplementary figure 8: Representative LCO HPLC elution profile 17
Supplementary figure 9: Representative LCO LC-MS spectra 18
Supplementary figure 10: Spinach germination 19

Supplementary figure 11: Corn germination experiments 20
Supplementary figure 12: Corn greenhouse 21

### Supplementary discussion

#### Comparison to cyanobacterial co-culture systems

As bioproduction technologies have expanded, co-culture and cross-feeding has been explored as a possible solution to lower feedstock costs while supporting the existing infrastructure of engineered heterotrophs. Efforts towards autotrophic-heterotrophic co-cultures have primarily focused on cyanobacteria as the autotroph[1,2]. Cyanobacteria are an obvious choice as they natively produce sucrose as an osmoprotectant—rather than a carbon source—to high concentrations without toxicity, making it an attractive feedstock-producer for heterotrophs. Engineered cyanobacterial strains able to convert and export up to 80% of their fixed carbon successfully fed three phylogenetically distinct heterotrophic microbes (*E. coli*, *B. subtilis*, and *S. cerevisiae*)[3]. However, cyanobacteria produce reactive oxygen species through photosynthesis and protective cyanotoxins, which are ultimately toxic to the heterotrophs. While cyanobacteria have higher solar-to-biomass conversion efficiencies than plants, efficiency remains 5-7% and is thermodynamically limited to ~12%—several fold lower than photovoltaics[4]. In addition to their biological limitations, there are a variety of implementation constraints that hinder industrial scale-up. Because cyanobacteria grown at scale require sunlight, two common culturing methods allow for optimal sunlight penetration: pools and photobioreactors. The large shallow pools can only be used in certain regions, are susceptible to environmental changes and contamination—and so it is difficult to maintain consistent batch-to-batch cultivation. In an effort to mitigate some of these issues, these pools can be modified to grow the cyanobacteria in small diameter tubing, but this kind of containment often deteriorates from radiation exposure as well as generates substantial plastic waste[5]. Because these issues are all challenges for cyanobacteria monoculture, it is not clear how a co-culture system would be successfully implemented at scale.

#### Sucrose production calculations

In comparing the different modes of sucrose production we calculated the respective productivities per hectare of land per year. Because the footprints of fermenters are orders of magnitude smaller than the land area needed to grow cyanobacteria and crops, we drew an equivalence to the land area needed to satisfy the $H_2$ demand were it to be derived from photovoltaics and water splitting. We compare sugarcane and engineered cyanobacteria[6] with our engineered *C. necator*. Solar-to-biomass conversion in plant crops have an annual efficiency of 1% and 3.5% (C3) or 4.3% (C4)[7,8], while cyanobacteria have an efficiency of 3% in open pools[9] and 5-7% in bubbled bioreactors[10]. For *C. necator*, the solar-$H_2$-efficiency of photovoltaic cells coupled to water electrolysis is 14%[4]. Land area was estimated based on an NREL case study[11] reporting 2 kWh $H_2$ $kg^{-1}$ and PVWatt calculator set to the Los Angeles area, we have 2,510,000 kWh $ha^{-1}$ $yr^{-1}$. Based on a California Energy Commission report, we assumed 3.3-3.6 g of biomass per gram of $H_2$[12]. Of the accumulated biomass, sugarcane is 20% sucrose, *S. elongatus* can be optimized to convert 80% of its biomass into sucrose, and this work (unoptimized) reports a biomass-to-sucrose conversion of 11.3%. Applying these values at hectare scale, we assume sugarcane produces 30-70 t $ha^{-1}$ $yr^{-1}$ biomass[13]; at a conversion of 20% that yields 14 t $ha^{-1}$ $yr^{-1}$ sucrose. Likewise, cyanobacteria in open-pond designs produce 25-50 t $ha^{-1}$ $yr^{-1}$ sucrose[14]; at a conversion of 80% that yields 40 t $ha^{-1}$ $yr^{-1}$ sucrose. Our engineered *C. necator*, in such a system, can reach a productivity of 510 t $ha^{-1}$ $yr^{-1}$ sucrose which is a 35-fold increase over sugarcane and 13-fold increase over cyanobacterial ponds.

#### Carbon footprint of plastics calculations

Coupled with its production, petrochemical plastics—depending on the type— persist in the environment for long periods of time that are largely undetermined (ranges from decades to thousands of years)[15]. Unfortunately, the most commonly used bioplastics (e.g., PLA, bio-PET) do not show much improvement in this context[16]. We propose that microbial production of biodegradable bioplastics through gas fermentation is a plausible alternative to reduce both GHG emissions and pollution.

We used the $CO_2$e values for the production, conversion, and end-of-life of PET, PP, PLA, and PHA made from sugarcane from Zheng et al. We calculated our potential $CO_2$e emissions from the energy input for a scaled system (personal correspondence) [17]. We assumed the same emissions for processing for $CO_2$-derived PHA as sugar-derived PHA. Biodegradability of these plastics in the environment is still not well characterized and depend on a variety of physical and chemical characteristics of the product itself (e.g., dimensions, compounding, blending)[18]. Because of this we use a qualitative approach to how these materials biodegrade naturally. Depending on the context, PET and PP can degrade into nanoplastics over decades or persist in the environment indefinitely. The functional outcomes of PLA in industrial composters remains unclear as the standard cycle times tend not to be long enough to degrade most PLA products, but they will degrade if the conditions are optimized[19]. PLA biodegradation in the environment is also unclear but appears to be limited to specific forms (e.g., agricultural mulch films). In contrast, PHA biodegradation in the environment is well established and can degrade within weeks to years depending on the product specification and environment[20–22].

#### Carbon footprint of fertilizer calculations

Aligned with our focus on industrial bioproduction to not compete for arable land, we sought to produce a plant growth enhancer that could be used to offset fertilizer use. Despite their central role in supporting modern agriculture, synthetic fertilizers generate significant GHG emissions, ecological hazards, and consequent adverse health and economic effects. As such ,more sustainable solutions are needed to plant growth and fertilization[30,31].

The average amount of $NH_3$ fertilizer applied in the United States is 92 kg $ha^{-1}$ $yr^{-1}$ [23,24]. The average $CO_2e$ from industrial Haber-Bosch process is 2.6 kg $CO_2$ $kg^{-1}$ of $NH_3$[25]. This amount of $NH_3$ generates 239.2 kg $CO_2$ $ha^{-1}$ $yr^{-1}$. We compare synthetic fertilizers to cover and inter-cropping and LCO addition. Cover cropping is a strategy used to improve the soil for the main crop by growing different plants when not growing the main crop. Intercropping is also used to improve the soil but does so by growing the main crop with others simultaneously, typically the additional crop is a legume. In our calculation, we included a variety of legumes and crops, locations, soil types, and application times[26–31]. A wide variety of responses were observed with cover/intercropping and there are many management considerations that must also be included when using this method of growth enhancement[32,33].

We assumed that the 40% increase conferred by synthetic fertilizer[28] in productivity is 100% of possible yield. The growth that results from intercropping or LCO addition was subtracted from the equivalent amount of fertilizer. That reduction is represented in the offset $CO_2e$ not generated from the alternative strategies. We applied these values to corn, whose production in the U.S. is approximately 11.1 tons $ha^{-1}$ (2019)[34]. We then determined the percent increase conferred by LCOs on different plants[34–37]. Two different scenarios were considered: 1) the growth enhancements in field studies (current) and 2) optimized conditions in greenhouses (potential). Because multiple studies addressing the same crop were limited, different crop species were used in calculations (e.g., corn, soybean, artichoke, rice, pea, and vetch[34–36,38]). We assumed no additional $CO_2$ drawdown from $CO_2$ fixation by *C. necator* to produce the LCOs.

### Supplementary figures

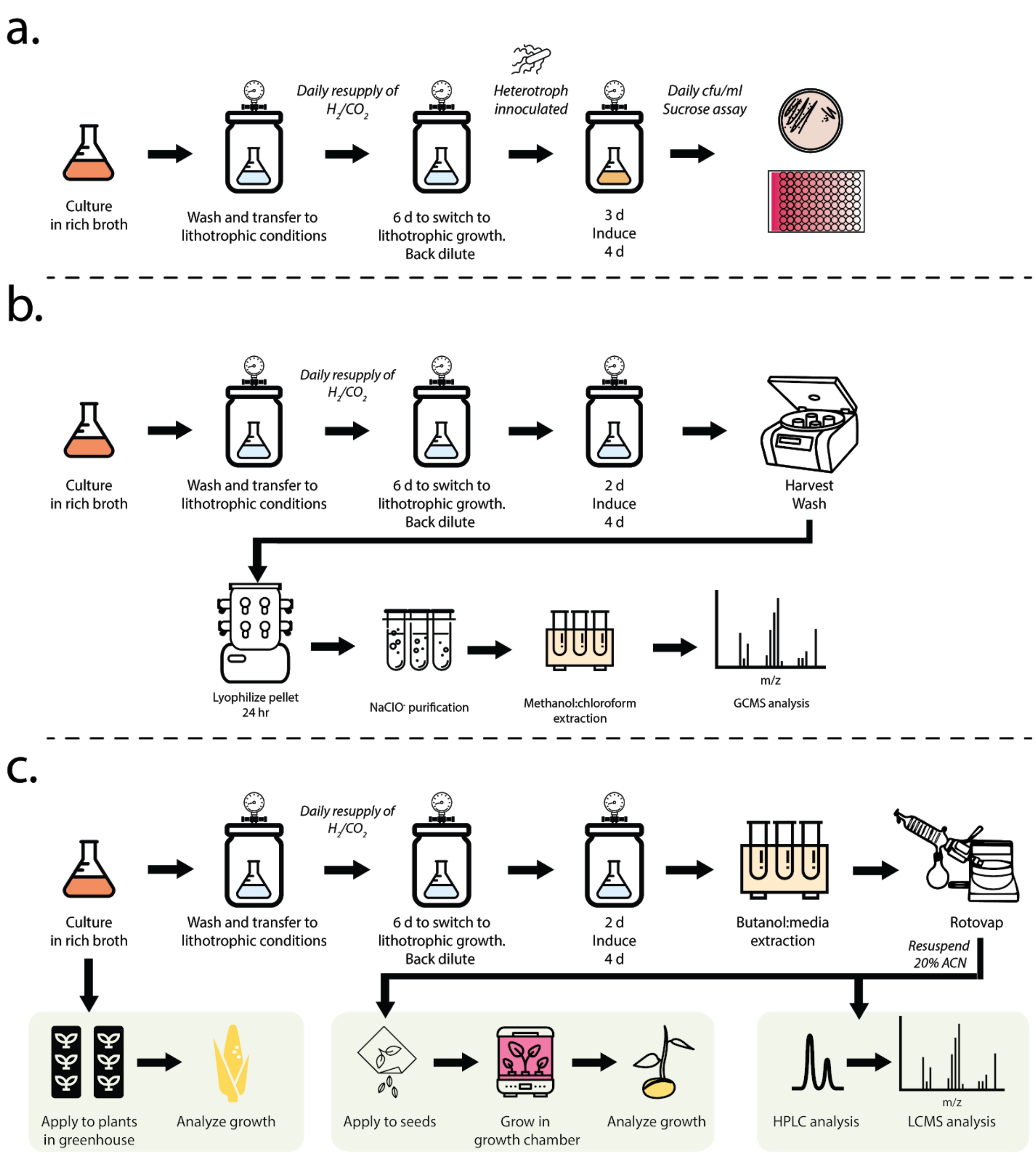

Supplementary figure 1: Experimental setup. All strains were initially inoculated in rich broth, washed, and inoculated in minimal Schuster media, placed in a vacuum jar, and supplied $CO_2$ and $H_2$ as the sole carbon and energy source, respectively. These cultures were then grown until they reach an $OD_{600}$ = 2-3, approximately 6 days. Once the cultures had switched to lithoautotrophic metabolism they were back-diluted into fresh media at $OD_{600}$ = 0.2 (for PHAs and LCOs) or $OD_{600}$ = 0.5 (for sucrose). After induction, the cultures were resupplied with fresh $CO_2$ and $H_2$ daily or every other day. **a.** Sucrose: Three days after back-dilution lithotrophic sucrose-producing *C.necator* were induced and inoculated with *E. coli* at $OD_{600}$ = 0.01. The co-culture was grown for an additional 7 days, plated and assayed for sucrose concentration every other day. **b.** PHAs: PHA-producing strains were induced at $OD_{600}$ = 1 or approximately 2 days in nitrogen-limiting media (if needed, acrylic acid was also added at this time). Strains were then grown for an additional 4 days to accumulate PHAs. Cells were then washed, lyophilized and lysed by $NaClO^-$. The PHA pellets were lyophilized, then subjected to methanolysis. 3-hydroxy acids were solubilized in chloroform and then analyzed by GC-MS. **c.** LCOs: LCO-producing strains were induced at $OD_{600}$ = 1 or approximately 2 days after back-dilution. After an additional 4 days of growth they were harvested, washed, subjected to butanol extraction, then concentrated by rotary evaporator. Samples to be analyzed by HPLC and LC-MS were solubilized in 20% acetonitrile. Samples to be applied for germination experiments were solubilized in water and applied to seeds. The seeds were grown in a growth chamber for 9 days and then analyzed. Samples that were applied to greenhouse experiments were purified from rich media due to volume limitations of the lithotrophic conditions (50 mL). Purified LCOs were applied to corn seeds, which were then planted. LCOs were applied a second time when planted. After 2 weeks the corn plant growth was analyzed.

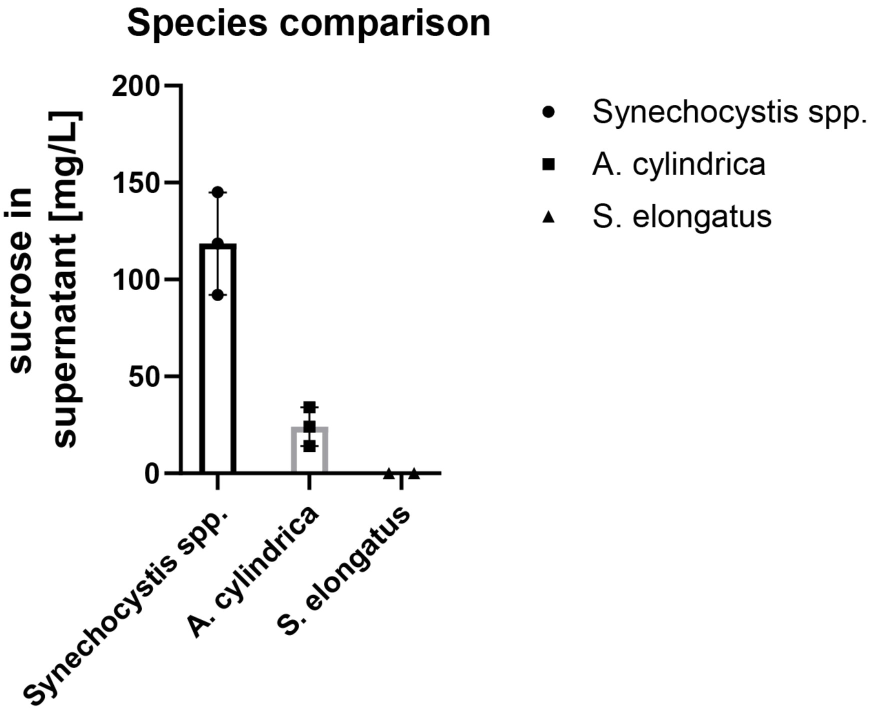


Supplementary figure 2. Comparison of sucrose producing enzymes from different cyanobacterial species expressed in *C. necator*. Sucrose phosphate synthetase and sucrose phosphate phosphatase from cyanobacterial species were expressed in *C. necator* and sucrose production in supernatant was determined after 7 days. Shown are three biological replicates with mean and standard deviation.

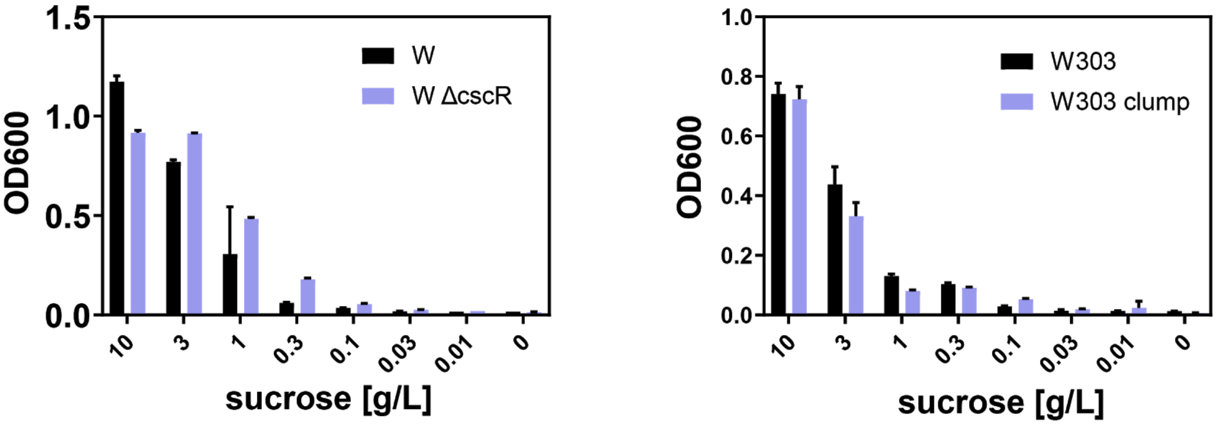


Supplementary figure 3 Sucrose titration. *E. coli* W and *S. cerevisiae* W303 strains were grown in Schuster media supplemented with varying concentrations of sucrose. $OD_{600}$ was recorded after 2 days of anaerobic growth. Reported are mean values of three biological replicates with error bars indicating standard deviation.

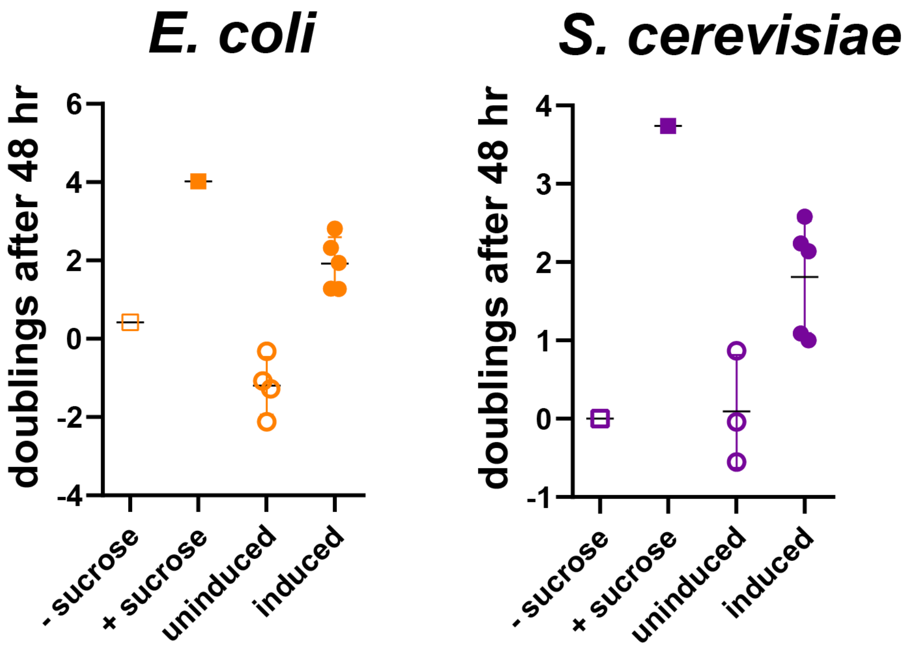


Supplementary figure 4. Heterotroph growth in *C. necator* supernatant. *E. coli* PAS842 and *S. cerevisiae* PAS844 were grown for 2 days anaerobically at 30 °C in supernatant from *C. necator* PAS837 that was grown lithotrophically for 7 days with and without induction. Colony count (cfu/mL) was assessed by plating at the beginning of the experiment and after 48 h and doublings were calculated. As a control, heterotrophs were grown in Schuster media with and without sucrose. In all conditions except for induced *C. necator* supernatant, 0.3% arabinose were added.

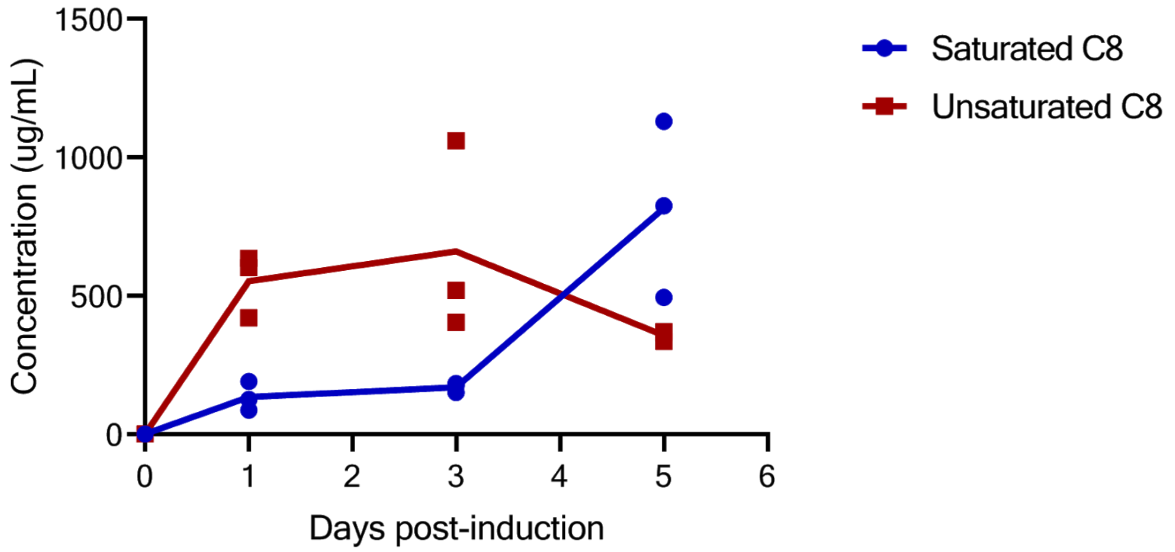


Supplementary figure 5: Octanoate production by *C. necator* expressing chim4 TE. Concentration as determined by GC-MS analysis. Known concentrations of C6, C8, C10, and C12 fatty acids were used to generate a standard curve and to quantify the production of single fatty acid species. Timepoints indicate days post-induction. Wildtype samples were below detection (data not shown).

| Strain | 828 | 828 | 829 | 829 | 830 | 830 |
|---|---|---|---|---|---|---|
| Acrylic acid | - | + | - | + | - | + |
| %DCW | **43.1±6.4** | **32.5±13.6** | **24.1±13.4** | **30.7±4.5** | **26.7±15.5** | **41.8±3.7** |
| **Strain** | **831** | **831** | **832** | **832** | **833** | **833** |
| Acrylic acid | - | + | - | + | - | + |
| %DCW | **36.9±8** | **36.2±8** | **54.4±9.9** | **45.5±0.01** | **23.2±17** | **33.5±7.2** |

Supplementary figure 6: PHA content relative to dry cell weight (DCW). Reported are mean values and standard deviation of three biological replicates for each strain.

a

| 3HA Chain length | PAS827 | PAS826 | PAS826 | PAS828 | PAS828 | PAS829 | PAS829 |
|---|---|---|---|---|---|---|---|
| C4 | 0±0 | 100±0 | 100±0 | 95.6±3.5 | 97.5±2.8 | 20.3±4.6 | 23.5±0.7 |
| C6 | 0±0 | 0±0 | 0±0 | 0.2±0.2 | 0.8±0.9 | 6.6±1.7 | 8.5±1.4 |
| C8 | 0±0 | 0±0 | 0±0 | 0.9±0.9 | 0±0 | 64.5±2.7 | 58.4±0.6 |
| C10 | 0±0 | 0±0 | 0±0 | 0.7±0.7 | 0±0 | 7.9±1.2 | 6.2±0.04 |
| C12 | 0±0 | 0±0 | 0±0 | 2.51±2.2 | 1.6±2.8 | 0.7±1.1 | 2.5±0.1 |
| C14 | 0±0 | 0±0 | 0±0 | 0±0 | 0±0 | 0.1±0.2 | 0.9±0.1 |
| Native phaC1$_{Cn}$ | - | + | + | + | + | - | - |
| Plasmid | - | - | - | + | + | + | + |
| Acrylic acid | - | - | + | - | + | - | + |

b

| 3HA Chain length | PAS827 | PAS826 | PAS826 | PAS830 | PAS830 | PAS831 | PAS831 |
|---|---|---|---|---|---|---|---|
| C4 | 0±0 | 100±0 | 100±0 | 41.5±1.2 | 27.4±4.5 | 21.3±1.3 | 18.9±3.4 |
| C6 | 0±0 | 0±0 | 0±0 | 4.2±3.1 | 7.2±2.3 | 5.5±6.4 | 8.8±1.5 |
| C8 | 0±0 | 0±0 | 0±0 | 48.9±5.9 | 54.6±0.7 | 62.6±7.6 | 61.4±1.7 |
| C10 | 0±0 | 0±0 | 0±0 | 4.1±3.5 | 6.7±2.5 | 7.2±1.0 | 7.1±0.3 |
| C12 | 0±0 | 0±0 | 0±0 | 1.3±1.2 | 2.1±0.6 | 1.8±1.6 | 2.6±0.1 |
| C14 | 0±0 | 0±0 | 0±0 | 0±0 | 2.1±1 | 1.3±2.3 | 1.2±1.3 |
| Native phaC1$_{Cn}$ | - | + | + | + | + | - | - |
| Plasmid | - | - | - | + | + | + | + |
| Acrylic acid | - | - | + | - | + | - | + |

c

| 3HA Chain length | PAS827 | PAS826 | PAS826 | PAS832 | PAS832 | PAS833 | PAS833 |
|---|---|---|---|---|---|---|---|
| C4 | 0±0 | 100±0 | 100±0 | 65.6±6 | 40.6±13.7 | 22.4±15.1 | 14.4±1.4 |
| C6 | 0±0 | 0±0 | 0±0 | 0.4±0.7 | 1.5±2 | 1.3±1.1 | 5.8±6.5 |
| C8 | 0±0 | 0±0 | 0±0 | 2.7±2.4 | 5.3±1.5 | 7.1±2.6 | 5.8±5.1 |
| C10 | 0±0 | 0±0 | 0±0 | 7.7±1.2 | 13.6±2.7 | 22.2±5.2 | 13.2±11.4 |
| C12 | 0±0 | 0±0 | 0±0 | 18.9±4.8 | 36.3±15 | 42.5±4.9 | 60.3±9.8 |
| C14 | 0±0 | 0±0 | 0±0 | 4.7±4.1 | 2.8±0.6 | 4.5±7.8 | 0.6±1 |
| Native phaC1$_{Cn}$ | - | + | + | + | + | - | - |
| Plasmid | - | - | - | + | + | + | + |
| Acrylic acid | - | - | + | - | + | - | + |

Supplementary figure 7: 3HA ratios in tailored PHAs. Values represent data in main text Fig. 3. **a.** PAS828 (phaC$_{Cn}$, pBAD *Uc*FatB2, phaC1$_{Pa}$); PAS829 (ΔphaC$_{Cn}$, pBAD *Uc*FatB2, phaC1$_{Pa}$). **b.** PAS830 (phaC$_{Cn}$, pBAD chim4, phaC1$_{Ps}$) PAS831 (ΔphaC$_{Cn}$, pBAD chim4, phaC1$_{Ps}$). **c.** PAS832 (phaC$_{Cn}$, pBAD *Uc*FatB2, phaC2$_{Pa}$) PAS833 (ΔphaC$_{Cn}$, pBAD *Uc*FatB2, phaC2$_{Pa}$). Reported are mean values and standard deviation of three biological replicates for each strain. Fatty acids represented by: C4, bright red; C6, light green; C8, dark red; C10, light blue; C12, royal blue; C14, teal.

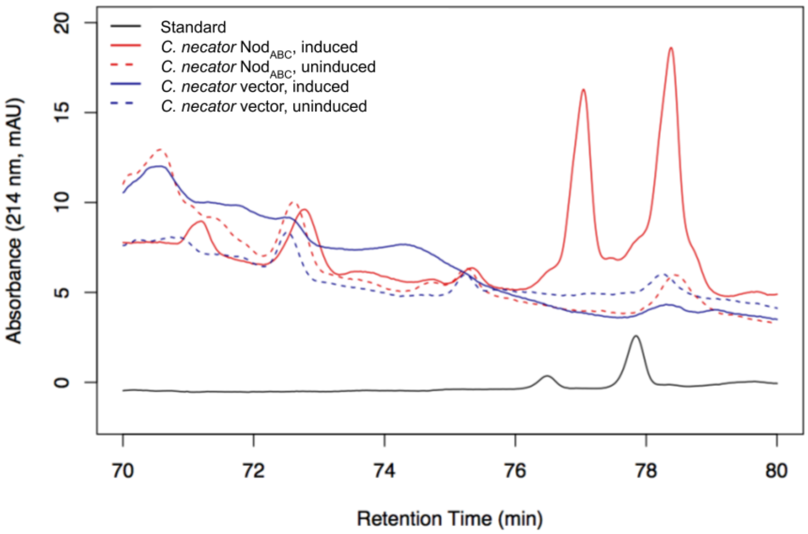


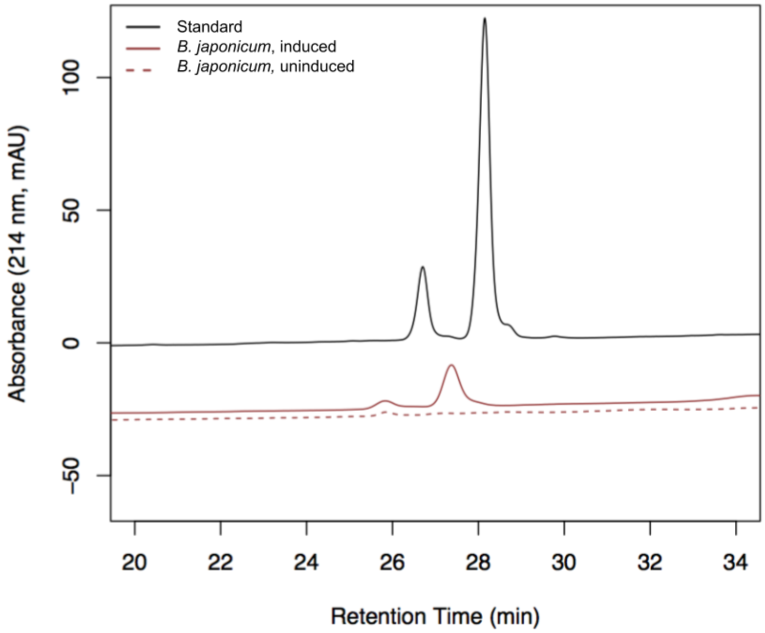


Supplementary figure 8**:** Representative LCO HPLC elution profile. (Top) Representative spectra from HPLC analysis of Nod Cn-V ($C_{18:1}$) (red). Purified extracts from induced and uninduced, vector control and engineered *C. necator*. (Bottom) Induced *B. japonicum* 100 compared to an LCO standard from *B. japonicum* 523C both indicate a characteristic double elution peak, which is seen in the engineered *C. necator* (PAS838) strain. Extracts from Standard (black), *B. japonicum* (brown), vector control *C. necator* (blue), and the engineered *C. necator* (PAS838) (red). Induced cultures are shown by solid lines and uninduced by dashed.

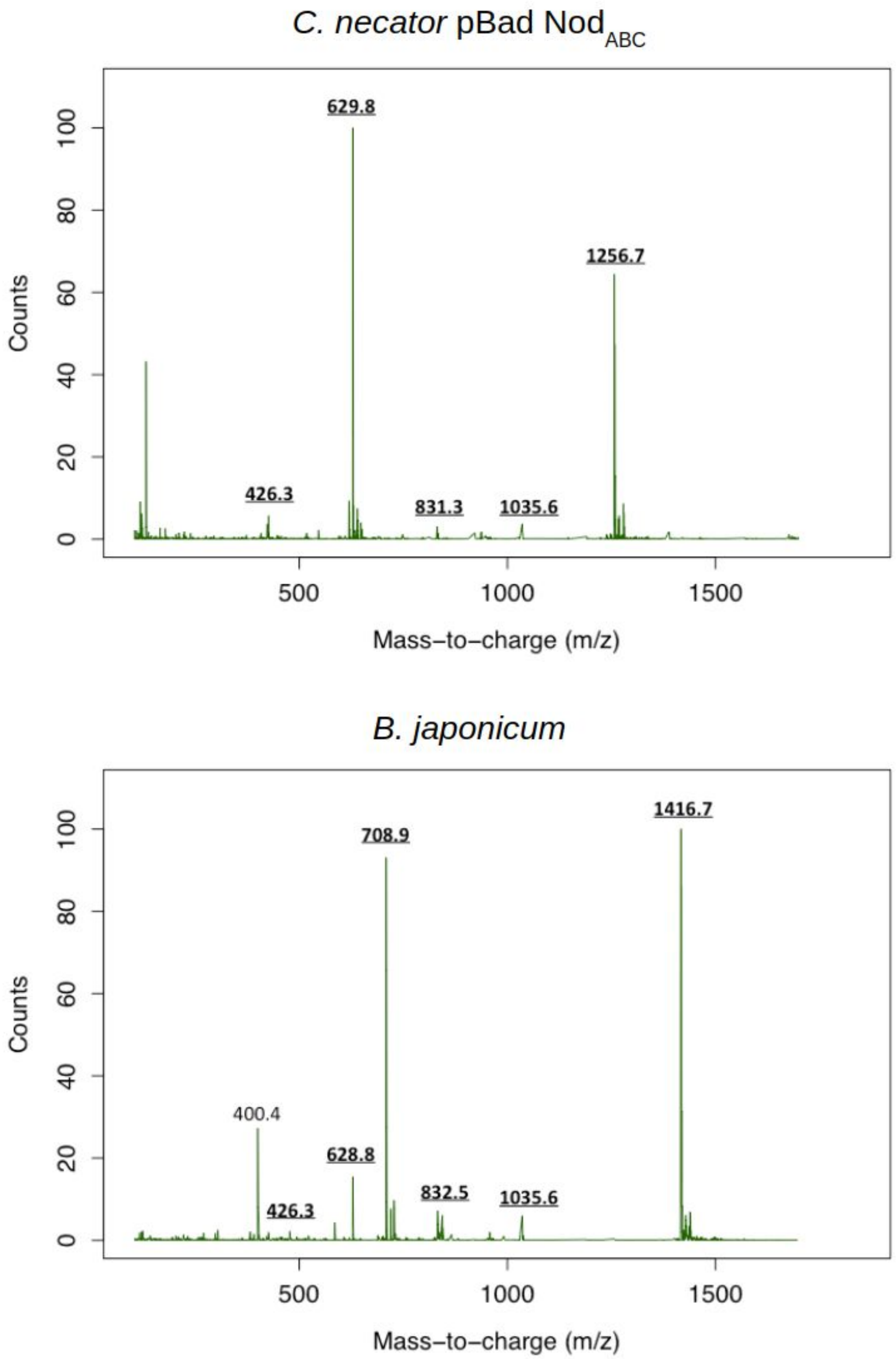


Supplementary figure 9**:** Representative LCO LC-MS spectra. (Top) Nod Cn-V ($C_{18:1}$) contains the characteristic peaks for the N-acetylglucosamine backbone with the largest peak at m/z = 1256 rather than m/z = 1416 indicating the lack of the fucose group found in *B. japonicum* 100. (Bottom) Nod Bj-V (C18:1 MeFuc). Relevant peaks are bold and underlined.

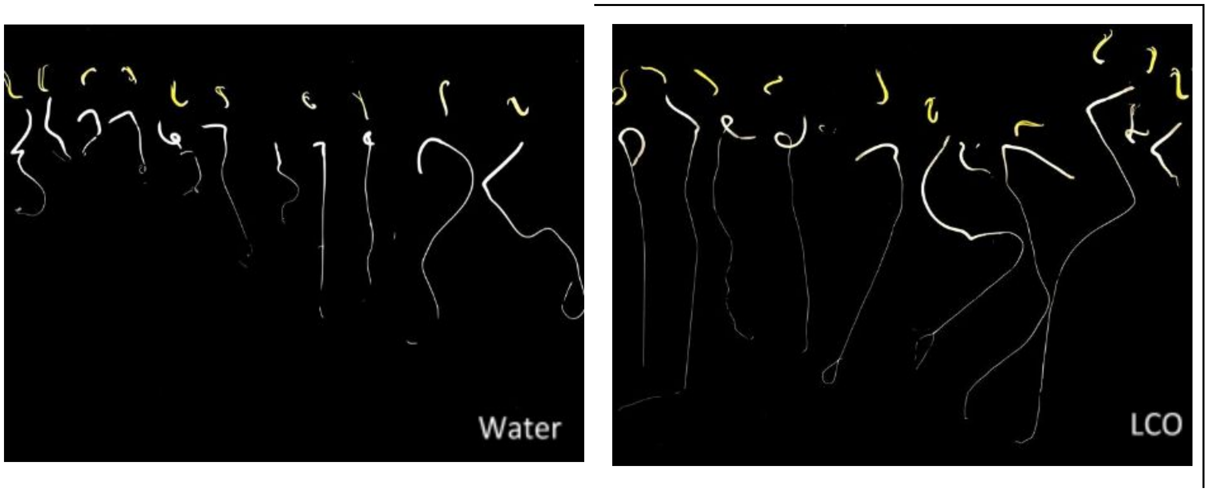


Supplementary figure 10**:** Representative germinated spinach seeds. The 10 longest seeds are shown in the water condition (left) and Nod Cn-V ($C_{18:1}$) condition (right).

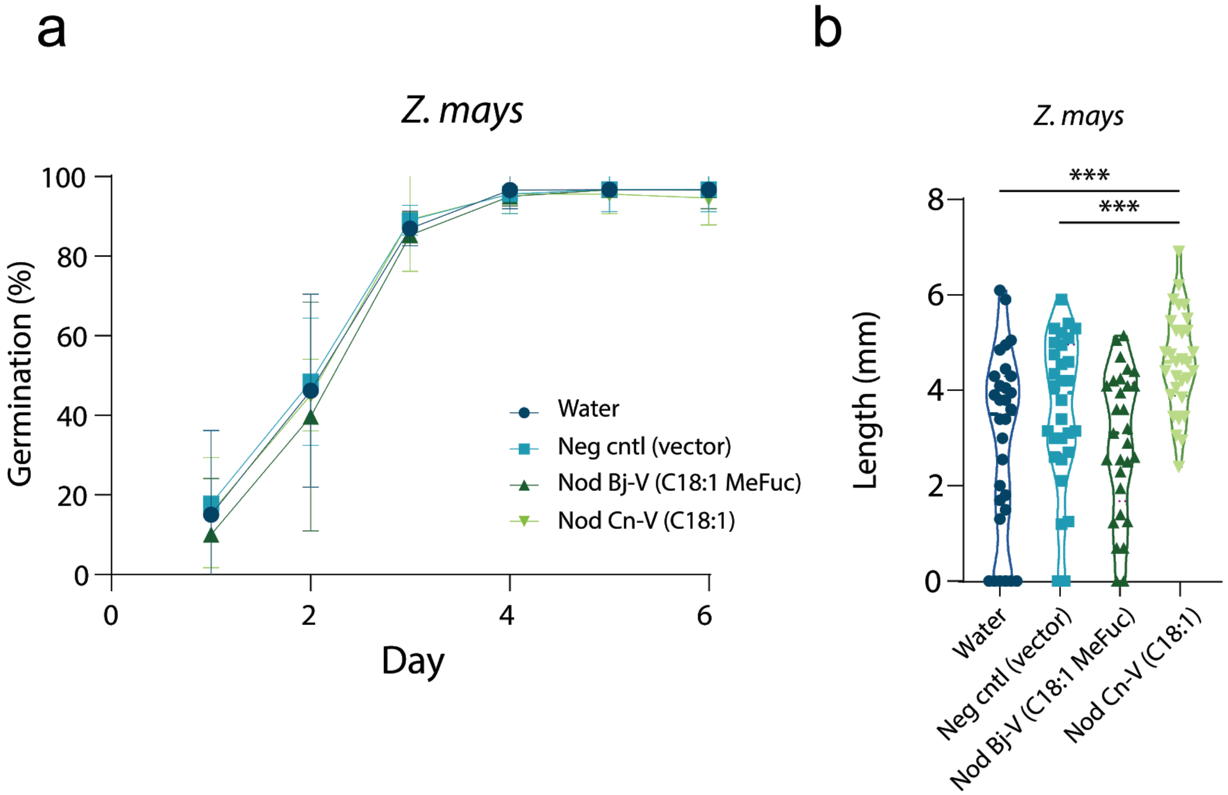


Supplementary figure 11: Corn germination experiments. (**a**) Germination rates in seeds in response to LCO application for corn. Seeds were treated with: water (dark blue circles), a vector control (blue squares), a standard LCO (Nod Bj-V (C18:1 MeFuc)) control from *B. japonicum* (dark green up-triangles) and the extract from *C. necator* vector control (bright down triangle). Corn shoot length showed increased length in the Nod Cn-V ($C_{18:1}$) compared to water (p=0.0003) and the vector control (p=0.0003). Asterisks indicate significance: *** <0.0001 as analyzed by multiple comparison one-way ANOVA with Tukey HSD post-hoc test.

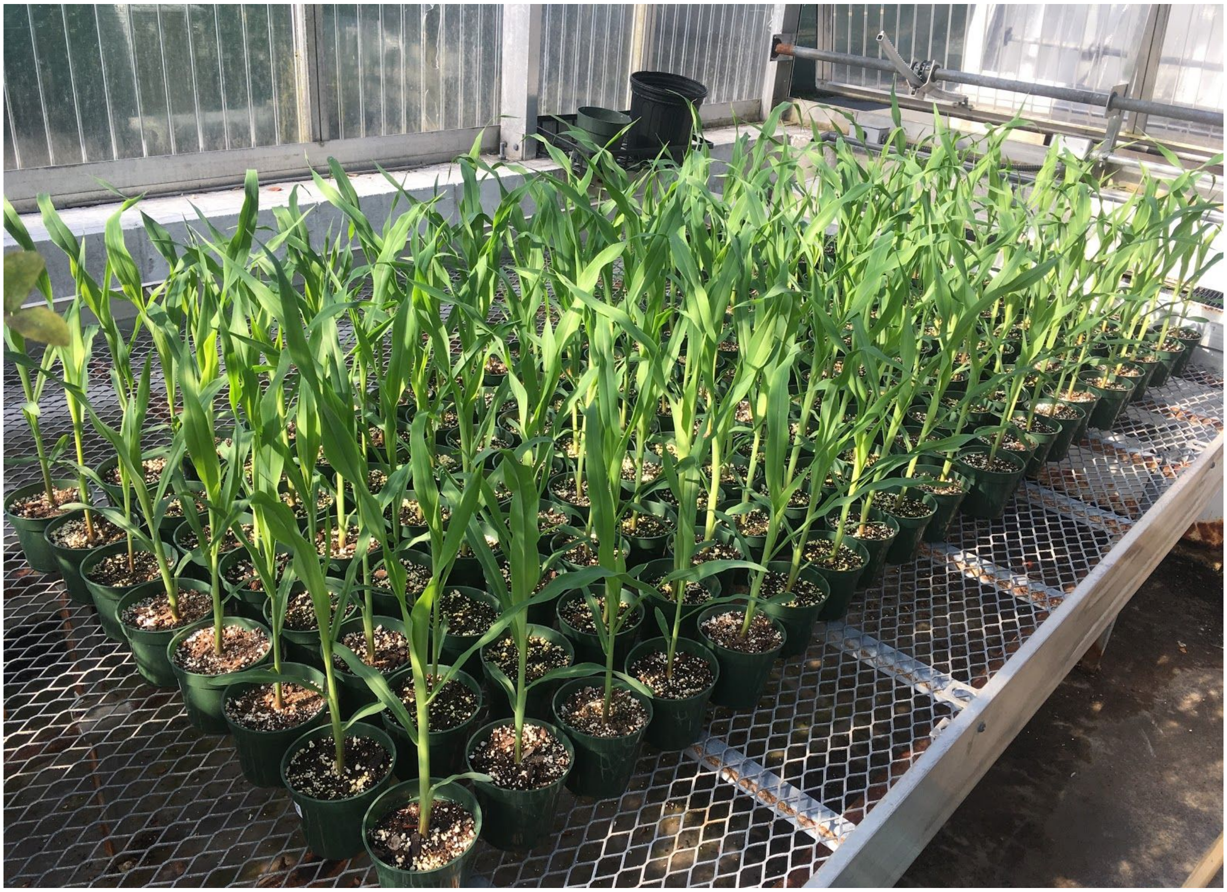

Supplementary figure 12: 160 corn plants were grown in a greenhouse for two weeks (10 replicates in each condition). Plants were grown and harvested in a blinded experimental setup.
